# Model biomolecular condensates have heterogeneous structure quantitatively dependent on the interaction profile of their constituent macromolecules

**DOI:** 10.1101/2022.03.25.485792

**Authors:** Julian C. Shillcock, Clément Lagisquet, Jérémy Alexandre, Laurent Vuillon, John H. Ipsen

## Abstract

Biomolecular condensates play numerous roles in cells by selectively concentrating client proteins while excluding others. These functions are likely to be sensitive to the spatial organization of the scaffold proteins forming the condensate. We use coarse-grained molecular simulations to show that model intrinsically-disordered proteins phase separate into a heterogeneous, structured fluid characterized by a well-defined length scale. The proteins are modelled as semi-flexible polymers with punctate, multifunctional binding sites in good solvent conditions. Their dense phase is highly solvated with a spatial structure that is more sensitive to the separation of the binding sites than their affinity. We introduce graph theoretic measures to show that the proteins are heterogeneously distributed throughout the dense phase, an effect that increases with increasing binding site number, and exhibit multi-timescale dynamics. The simulations predict that the structure of the dense phase is modulated by the location and affinity of binding sites distant from the termini of the proteins, while sites near the termini more strongly affect its phase behaviour. The relations uncovered between the arrangement of weak interaction sites on disordered proteins and the material properties of their dense phase can be experimentally tested to give insight into the biophysical properties and rational design of biomolecular condensates.

## Introduction

The phase separation of intrinsically-disordered proteins (IDP) into biomolecular condensates has taken centre stage in cellular physiology.(1) Biomolecular condensates (BC) appear in numerous locations in cells where they carry out many biochemical functions.(2, 3) They modulate enzymatic activity,(4-7), a function that has been reproduced with rationally designed peptides;(8-10) buffer protein concentration,(11) reduce noise in gene expression,(12) and regulate cell migration.(13) Phase separation of IDPs is crucial for the healthy functioning of neuronal synapses in the presynaptic axon(14, 15) and postsynaptic dendrite.(16, 17) However, dysfunctional phase separation underlies many pathological processes. Many neurodegenerative diseases involve aberrant phase transitions of IDPs,(18) and interfering with such transitions presages new routes for therapeutic advances in neurology.(19) Viral replication occurs within phase-separated inclusion bodies,(20) and proteins responsible for packing the RNA of SARS-Cov2 also undergo phase separation.(21) These processes depend on the ease with which enzymes and reactants can diffuse, fluctuate, and interact within.(22, 23) A better understanding of how the molecular architecture of scaffold IDPs influences the material properties of BCs would help elucidate the physical mechanisms underlying their functions and guide the identification of targets for drugs,(24) opening up new approaches to attack cancer and other diseases.(25-28)

Although many biomolecular condensates are fluid,(29) they do not behave as simple liquids. A slow transition from the fluid phase to a rigid, fibrillous phase has been observed *in vivo* and *in vitro* for IDPs including Huntingtin(30, 31), FUS,(32, 33) and alpha synuclein(18). Chromatin has been shown to undergo phase separation *in vitro*,(34) and to be mechanically restructured in a process that depends on its viscoelastic properties.(35, 36) Such experiments show that recapitulating the dynamics of condensates does not require the complex environment of a living cell.(23) It has been conjectured that *reversible* phase separation is indicative of cellular health while *irreversible* rigidification of BCs marks a cell’s transition into disease states.(29) Empirically classifying biomolecular condensates on a spectrum from fluid (healthy) to rigid (disease) motivates us to learn how their material properties arise from their constituent IDPs.(37-39) But although an enormous amount of experimental data is accessible in online databases,(40-42) via web-based interfaces (https://forti.shinyapps.io/mlos/),(43) there is no mechanistic understanding of how IDP molecular architecture controls the structure of their dense phase.

Computational modelling has been extensively used to connect molecular details such as multivalency and hydrophobicity to IDP phase behaviour in model systems.(44-47) Exquisite detail on residue-residue interactions(48) and the conformational dynamics of IDPs within condensates is revealed by Atomistic Molecular dynamics (aaMD) simulations,(23, 49) but these are limited to near-molecular system sizes and short times. Coarse-grained simulation techniques simplify the atomic details of proteins in order to simulate thousands of molecules.(50-55) A minimal model of the phase separation of multivalent proteins is the patchy particle scheme in which proteins are treated as spheres with attractive sites on their surface.(56, 57) These models, however, ignore conformational fluctuations that may be important for flexible proteins. Other coarse-grained models represent an IDP as a linear polymer of monomers (or *beads*) held together by bonds (or *springs*). A bead represents one or more amino acids and the models differ chiefly in which atomic details are kept.(37, 44, 45, 50, 53, 58) The translational and conformational degrees of freedom of the IDPs are therefore retained whilst their enthalpic interactions are simplified to increase the accessible length and time scales. Recent reviews describe how modeling is able to reveal physical principles underlying phase separation of IDPs.(51, 59)

In this work, we go beyond establishing the phase boundaries of a model IDP in aqueous solution, and quantitatively study its dense phase. An IDP is represented as a semi-flexible polymer with multiple, punctate, attractive sites immersed in a *good* solvent.(4, 60) Our results are thus distinct from previous applications of similar *stickers and spacers* models, in which the polymers are in a poor solvent and possess a large fraction of hydrophobic monomers.(37, 38, 45, 61, 62) Although motivated by the study of IDPs, our model is also applicable to the phase separation of uncharged, associative polymers.(63) We use the coarse-grained simulation technique of dissipative particle dynamics,(64, 65) which is suitable for simulating (uncharged) IDPs because of their polymeric nature and transient, non-specific interactions.(66) We find that the dense phase has a low (∼ mM) internal concentration and approximately 70% solvent by volume, both results in good agreement with experiments on the uncharged IDP FUS.(33, 67) The IDPs form a structured fluid network in which their binding sites transiently meet at junctions. The length scale between the junctions is much larger than the monomer size, and varies more strongly with the binding site separation than with their affinity. It is reminiscent of the diffraction peaks observed in SANS experiments on the dense phase of the multivalent nucleolar protein NPM1.(68) The spatial heterogeneity is revealed further using graph theoretic measures,(69, 70) which have been applied to metabolic networks,(71) allosteric pathways,(72) and the importance of residue mutations in proteins.(73) We map the junctions to the nodes of a graph, and place edges between nodes spanned by at least one polymer. The local clustering coefficient quantifies the crowding of polymers around the nodes.(74, 75) Thirdly, the dense phase evolves on multiple timescales, indicating a complex internal dynamics of the IDPs.(23) Finally, our model predicts that the location of attractive domains on IDPs is critical for their phase separation. When the binding sites at the polymer endcaps are disabled, the dense phase dissolves.

## Methods

### Dissipative particle dynamics simulation technique

We use the Dissipative Particle Dynamics simulation technique (DPD) to study the phase behaviour of a series of model IDPs. The source code for the simulations is available on Github.(76) DPD is a coarse-grained, explicit-solvent simulation technique invented to study the hydrodynamic behaviour of complex fluids.(64, 65) It has since been applied to many soft matter systems including amphiphilic membranes,(77-79) vesicle fusion,(80, 81) and domain formation in vesicles(82), among many others.(83) It is highly suited to simulations of (uncharged) IDPs because the interactions are weak, and it has been shown that polymers in DPD exhibit self-avoiding walk scaling.(84) DPD is able to follow the evolution of fluid systems over large length and time scales by grouping atoms or atomic groups into beads, and replacing complex, interatomic potentials by effective forces that are softer and short-ranged. This reduces the number of degrees of freedom being integrated, and allows a larger integration time step in the equations of motion.

Beads in DPD have mass *m* and interact via three non-bonded interactions that are soft, short-ranged (vanish beyond a fixed length-scale *d*_0_), pairwise additive, and conserve linear momentum. A conservative force gives each bead an identity such as hydrophilic or hydrophobic:

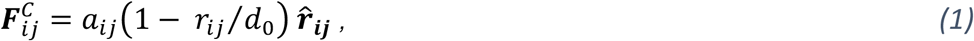

for *r*_*ij*_ < *d*_0_. The maximum value of the force is *a*_*ij*_; ***r***_***ij***_ = ***r***_***i***_ − ***r***_***j***_ is the relative position vector from bead j to bead i, *r*_*ij*_ is its magnitude, and 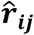 is the unit vector directed from bead *j* to bead *i*. The other two non-bonded forces constitute a thermostat that ensures the equilibrium states of the simulation are Boltzmann distributed.(65) The dissipative force is:

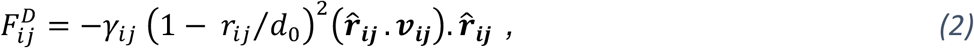

where *γ*_*ij*_ is the strength of the dissipative force and ***ν***_***ij***_ is the relative velocity between beads *i* and *j*. This force destroys relative momentum between interacting particles. The random force is:

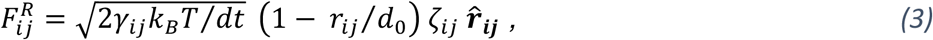

where *k*_*B*_*T* is the system temperature and *ζ*_*ij*_ is a symmetric, uniform, unit random variable that is sampled for each pair of interacting beads and satisfies *ζ*_*ij*_ = *ζ*_*ji*_, ⟨*ζ*_*ij*_(*t*)⟩ = 0 and ⟨*ζ*_*ij*_(*t*)*ζ*_*kl*_(*t*′)⟩ = (*δ*_*ik*_*δ*_*jl*_ + *δ*_*il*_*δ*_*jk*_)*δ*(*t* − *t*′). This force creates relative momentum between pairs of interacting particles *i* and *j*. The factor 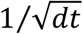 is required in the random force so that the discretized form of the Langevin equation is well defined.(65)

Molecules are constructed by tying beads together using Hookean springs with potential energy:

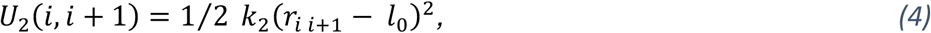

where the spring constant, *k*_2_, and unstretched length, *l*_0_ may be different for each bead type pair but are here fixed at the values 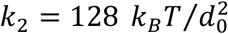 and *l*_0_ = 0.5 *d*_0_. The semi-flexible nature of IDPs is represented by a chain bending potential applied to the angle θ defined by adjacent backbone bead triples (BBB):

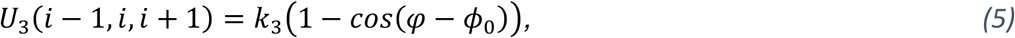

with parameters *k*_3_ = 5 *k*_*B*_*T* and *φ*_0_ = 0. All bonded and non-bonded interaction parameters are given in Table 1. For further details of the force field and DPD method applied to IDPs, the reader is referred to previous work.(47)

**Table 1.** Bead-bead conservative force parameters *a*_*ij*_ (in units of *k*_*B*_*T*/*d*_0_) and dissipative force parameters *γ*_*ij*_ (in units of 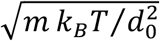) for all bead pairs, and Hookean spring potential parameters (in units of 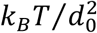 and *d*_0_ respectively). Note that E represents the end-cap beads and F the internal binding site beads for computational and visualisation purposes, but these bead types have identical interactions except when one or the other is turned off as described in the text. NA = not applicable. A three-body bending potential is applied to adjacent trips of backbone beads (BBB) with parameters *k*_3_ = 5 *k*_*B*_*T* and *φ*_0_ = 0.

| Bead Pairs | $a_{ij}$ | $\gamma_{ij}$ | $k_2$ | $l_0$ |
| --- | --- | --- | --- | --- |
| WW, WE, WF, | 25 | 4.5 | NA | NA |
| WB | 23 | 4.5 | NA | NA |
| BB, BE, BF | 25 | 4.5 | 128 | 0.5 |
| EE, EF, FF | Varies | 4.5 | 128 | 0.5 |

An IDP is represented as a linear polymer of hydrophilic backbone beads (B) interspersed by short segments of hydrophilic *sticky* beads (see Figure 1). Each segment contains 4 beads of type F that adopt approximately spherical binding sites. The endcaps are represented by 4 beads of type E. Bead types E and F have identical interactions except when one or other is disabled to represent disabling a binding site. They are coloured differently for distinctness only. Backbone beads are green, endcaps are red, and internal binding sites are yellow. The backbone beads are slightly more hydrophilic than binding sites to ensure they remain solvated in the dense phase. The solvent is represented by a single bead W and is invisible in all snapshots/movies for clarity.

**Figure 1.**
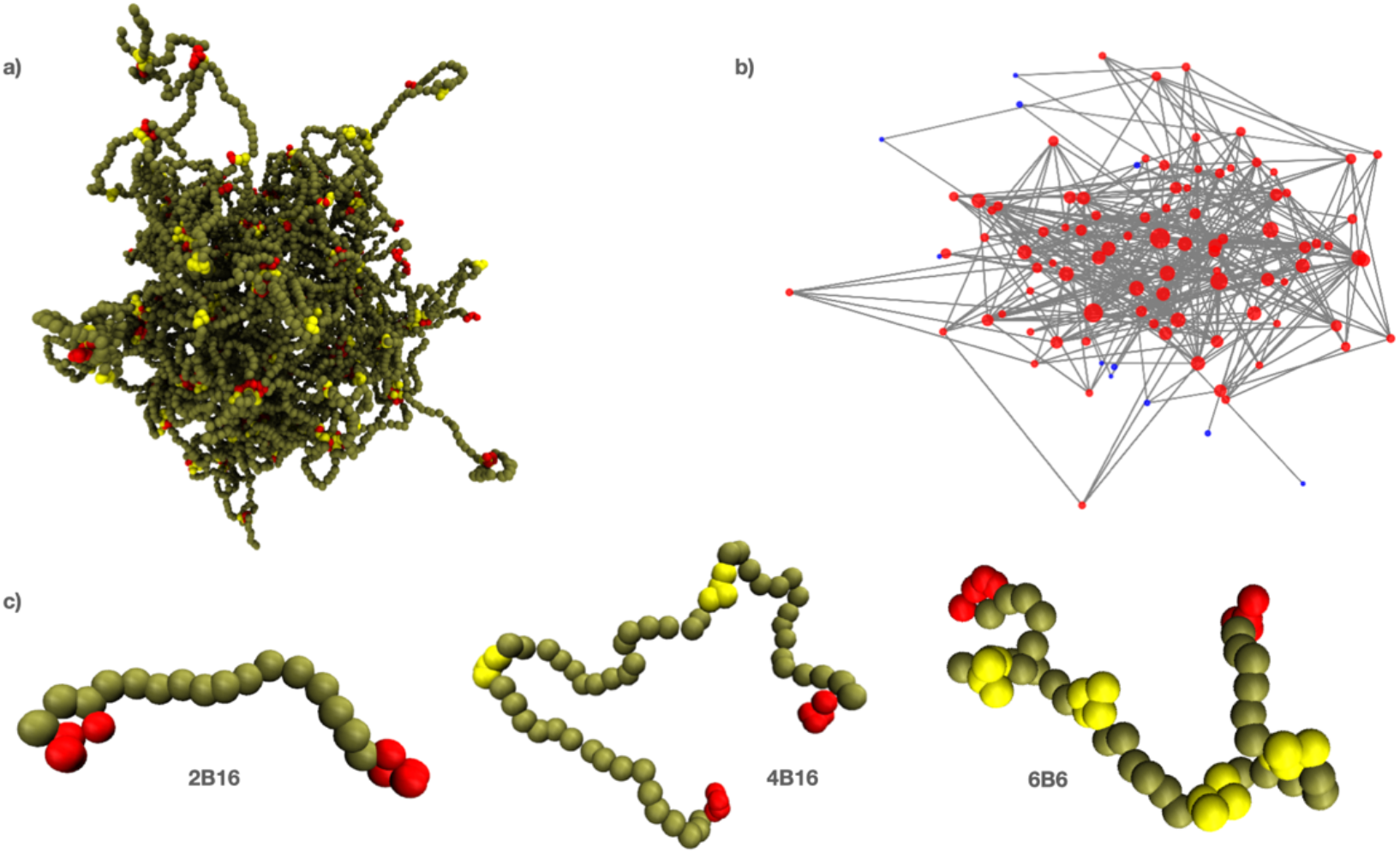
(**a**) Snapshot of 160 polymers of type 4B16 in solvent (invisible for clarity). (**b**) The equivalent graph for the largest connected component of the system shown in (a). Nodes of the graph are placed where the polymer binding sites touch and edges connect nodes spanned by at least one polymer. (**c**) Nomenclature for polymers with multiple binding sites (see Methods).

The number, location and affinity of the binding sites are the main parameters of the model. We use the nomenclature **nBm** (or **nIm**) to identify the structure of a polymer in the text, where **n** is the total number of binding sites (internal plus endcaps), and **m** is the number of backbone beads between adjacent binding sites (referred to as the *gap* between binding sites.) The letter **B** indicates that endcaps are present in the polymer and the letter **I** indicates that only internal binding sites are present. Selected binding sites can be turned on or off thereby changing the effective separation of the remaining active sites in a manner analogous to PTMs on a protein revealing/occluding specific interaction motifs. A typical phase separated droplet is shown in Figure 1a together with its equivalent graph in Figure 1b, which is further discussed in Section E. We refer to the dense phase equivalently as a droplet, network, or simply the dense phase. Three examples of polymers with different numbers of binding sites are shown in Figure 1c.

Simulations take place in a cubical box of size (48 *d*_0_)^3^ unless otherwise noted, and the system temperature is *k*_*B*_*T* = 1. A given number of IDPs are randomly distributed throughout the simulation box together with solvent particles to the constant density 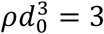. The system is evolved by integrating Newton’s laws of motion for all beads for at least 3 million time steps. The integration step size is 0.02 *τ*, where 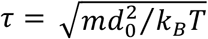 is the DPD timescale. The first million steps are discarded to allow the system to equilibrate, and ensemble averages are constructed by sampling from the remaining time. Because we are interested in equilibrium properties, we do not try to characterize the time-scale more precisely. One simulation of this system (3 million steps) requires 20 cpu-days on a single core of a 4.5 GHz Ryzen Threadripper 3970X machine. Further details of the DPD technique are given in the literature.(65, 83)

## Results

### A Dense phase concentration is modulated by binding site separation and affinity

We first explore how the phase behaviour of the model IDPs depends on the number and strength (or affinity) of their binding sites. These sites represent residue motifs with weak, uncharged, attractive interactions.(4, 60) Because the model IDPs are hydrophilic, the only driving force for phase separation comes from the attraction between these sites. The conservative interaction between beads of type *i, j* is characterized in DPD by a parameter *a*_*ij*_. We parametrize the binding site attraction by a dimensionless parameter *ϵ* that is defined in terms of the DPD conservative force parameters for the binding site/solvent beads (*a*_*EW*_ = *a*_*FW*_) and binding site/binding site beads (*a*_*EE*_ = *a*_*FF*_) via *ϵ* = (*a*_*EW*_ − *a*_*EE*_)/*a*_*EW*_. The value *ϵ* = 0 corresponds to no attraction, and *ϵ* ∼ 1 is a strong attraction between binding site beads.

Although representing an IDP as a semi-flexible polymer with discrete binding sites is a great simplification, the resulting model still possesses a large parameter space. At a minimum, the polymer length, bending stiffness, concentration, and the number, location and affinity of the binding sites must be specified. After a preliminary exploration, we focused on B48 polymers with 6 binding sites separated by 6, 8, or 10 backbone beads (referred to hereafter as 6B6, 6B8, 6B10 respectively). The top row of Figure 2 shows the dense phase of 6B6 polymers for three values of the affinity decreasing from left to right. The droplet shows little visible change in cohesion over this range. By contrast, the bottom row shows that a droplet composed of 6B10 polymers begins to dissolve for the same change in affinity. The stability of the dense phase is clearly sensitive to the affinity and separation of the binding sites: more widely-separated binding sites require a higher affinity to drive their phase separation.

**Figure 2.**
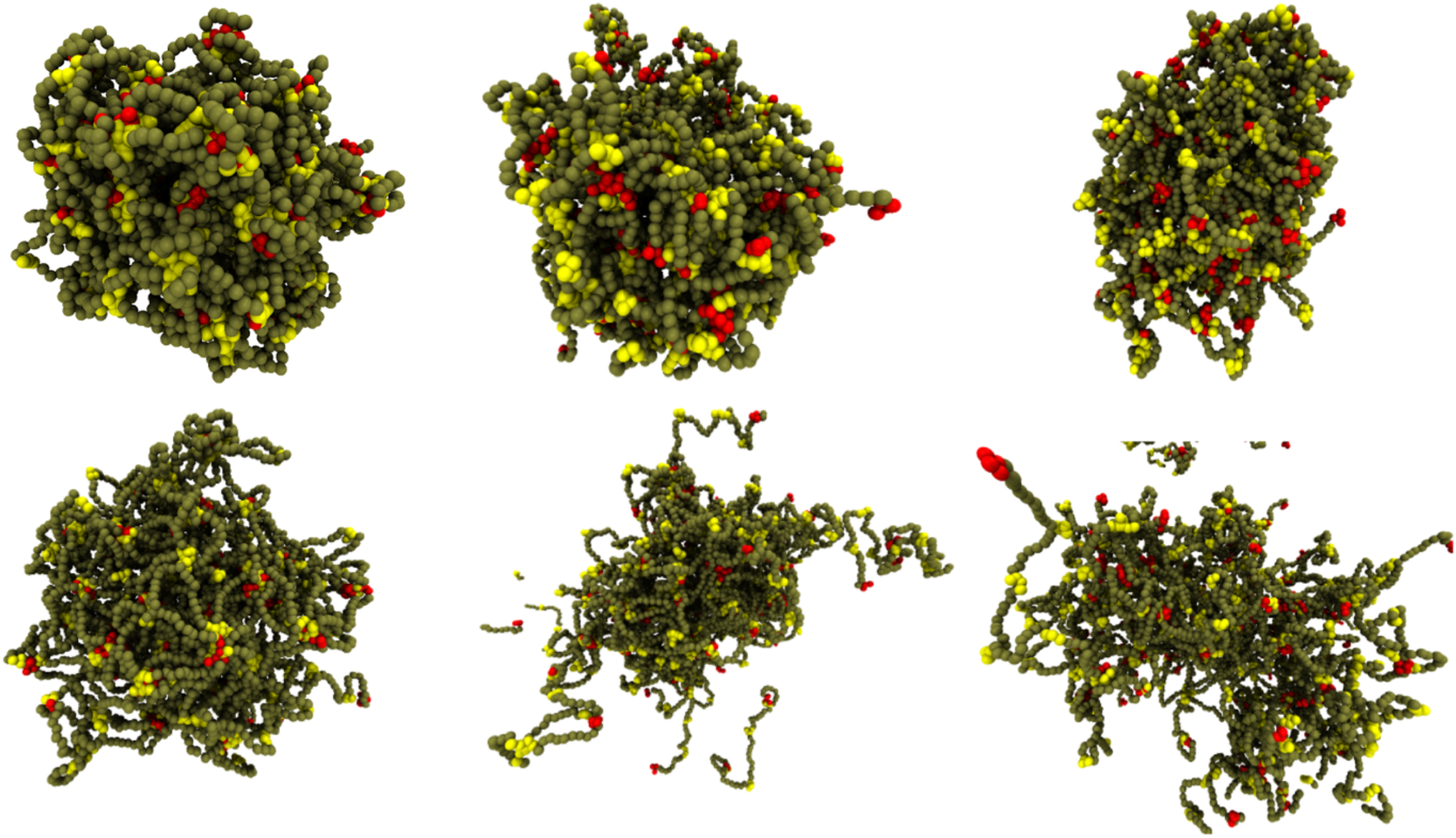
The dense phase stability depends on the binding site separation when their affinity is reduced. The 6B6 droplet remains phase separated on reducing the affinity, but the 6B10 droplet starts to dissolve. (Top row, left to right) 129 polymers 6B6 with affinity ε = 0.84, 0.76, 0.74; (Bottom row, left to right) 128 polymers 6B10 with the same affinities.

We establish a correspondence between the simulated droplets and experiments by calculating the concentration of the dense phase. Although it is difficult to measure the volume of arbitrarily-shaped droplets, their fluidity usually results in their being approximately spherical (cp. Fig 2), which enables their volume to be estimated from their radius of gyration. This is obtained from the coordinates of the polymers’ binding site beads under the assumption (verified by visual inspection of snapshots) that they are uniformly distributed throughout the droplet’s volume.

In equilibrium, the dense phase concentration C is:

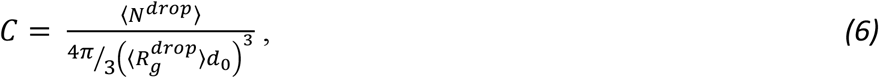

where ⟨*N*^*drop*^⟩ and 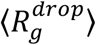 are the time-averaged number of polymers in the droplet and the droplet’s (dimensionless) radius of gyration respectively. The parameter *d*_0_ is the DPD length scale (see Methods section). To proceed further we have to map the DPD polymers onto an experimental IDP.

A common target for experimental investigation is the RNA-binding protein Fused in Sarcoma (FUS) that is implicated in the neurodegenerative disease ALS.(85) The low complexity, N-terminal domain of FUS (FUS-LC, residues 1-163) is enriched in three polar amino acids (Q, S, Y), has only two charged residues (D), and around one quarter hydrophobic residues (mainly G and P) and none of the most hydrophobic residues (I, F, L, M V). FUS-LC phase separates into fluid droplets(32) with a high viscosity.(33) It makes multiple, transient, weak interactions within the dense phase indicating that it has little or no secondary structure.(66, 86) The near absence of charge and secondary structure make it suitable for coarse-grained simulations.

We do not attempt to map the specific residue sequence of FUS-LC onto the coarse-grained polymers in our model. Instead, we assign an effective attraction to the punctate binding sites along the polymers thereby allowing a variety of arrangements of binding sites to be mapped onto the experimental FUS-LC system. Next, we choose the level of coarse-graining to connect the simulated IDPs with the FUS-LC protein. The DPD length scale *d*_0_ is fixed by equating the experimentally derived radius of gyration of FUS-LC with the value obtained from a simulation of a single polymer in the dilute phase: 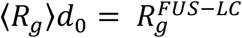. Figure S9 in the Supplementary Material shows the radius of gyration of each IDP studied obtained from independent simulations of a single polymer in dilute solution. It is evident that the model IDPs exhibit the conformational fluctuations of a self-avoiding walk for all the binding site affinities and polymer lengths studied.

Tomasso *et al*.,(87) and Marsh and Forman-Kay(88) have empirically fit the hydrodynamic radius *R*_*h*_ of a wide range of uncharged IDPs with the formula:

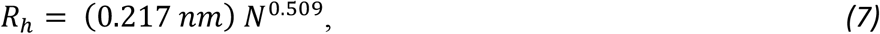

where N is the number of residues in the proteins, and the exponent is close to the Flory exponent of 1/2 for ideal chains.(89) In order to relate the hydrodynamic radius of an IDP to its radius of gyration one has to adopt a particular polymer model, e.g., treating it as an ideal chain or self-avoiding walk (SAW). Dünweg *et al*. used computer simulations to show that the hydrodynamic radius (*R*_*h*_) of a fluctuating polymer is related to its radius of gyration (*R*_*g*_) by *R*_*g*_/*R*_*h*_ = 1.5045 (ideal chain) or *R*_*g*_/*R*_*h*_ = 1.591 (SAW).(90) The radius of gyration of an ideal chain is related to its mean end-to-end length by 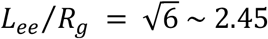, and Dunweg *et al*. further established that the corresponding relation for a self-avoiding walk is 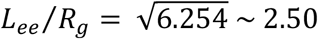. It is interesting to note that these relations imply that an IDP’s hydrodynamic radius is *smaller* than its radius of gyration, which is the opposite of a uniformly-dense sphere, for which the relationship is 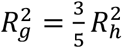. This implies that a fluctuating polymer in the dilute phase diffuses *faster* than a sphere with the same radius of gyration.

FUS-LC contains 163 residues and its hydrodynamic radius is predicted to be 2.9 nm from Eq. 2. Its radius of gyration is *R*_*g*_ = 4.36 *nm* using the relation for ideal chains or 4.61 nm for a SAW. The value of *d*_0_ is then fixed by: 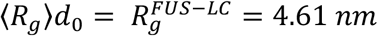. Combining all the numerical factors together with 1000/0.6 to convert the concentration from polymers/nm^3^ into mM in Eq. 1 allows the dense phase concentration to be determined from the formula 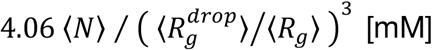, where we have adopted the SAW model for the polymer. The quantities in angle brackets are obtained by sampling from the equilibrated stage of the simulation. If the polymers are regarded as ideal chains instead of self-avoiding walks the numerical constant 4.06 changes to 4.8, corresponding to an increase in concentration of about 18% independent of the droplet size, which does not change our results significantly.

We find that the dense phase concentration lies in the range 1 - 15 mM for all systems with 4, 5, or 6 binding sites separated by 4 – 10 backbone beads, and for affinities between *ε* = 0.76 − 0.84 (Tables S2 and S3.) These values are in excellent agreement with recent experiments that find the dense phase concentration to be 2 mM for full length FUS,(67) and 7 mM(66) and 27.8 mM(33) for FUS-LC. We also show in Section 6 of the Supplementary Material that the dense phase contains 65-70% water by volume, a result also in good agreement with experiment.(33)

### B The spatial structure of the dense phase varies with binding site separation

We next determine how the spatial organisation of the model IDPs depends on the affinity and location of their binding sites. Close examination of Figure 2 shows that the binding sites (yellow for internal and red for endcaps) meet at junctions within the condensed phase that are separated by solvent-filled voids (Supplementary Movies M1-M5 show the equilibrium fluctuations of droplets composed of 6B6 and 6B10 polymers with affinities *ε* = 0.84, 0.74, 0.68). The distribution of binding sites at these junctions is highly heterogeneous. We quantify this observation by counting the number of binding sites on each junction (the junction *mass*) and measure the distance between them (the junction separation). Figure 3 shows how these observables vary over a wide range of the total polymer concentration. Because the droplet is a fluid, there is a continuous exchange of polymers between it and the surrounding dilute phase, whose rate depends on the binding site affinity. For this analysis, we identify the dense phase with the Largest Equilibrium Network (LEN), which is defined as the largest set of polymers that are connected into a network by at least two binding sites. This excludes polymers in the dilute phase and those dangling off the droplet by a single binding site.

**Figure 3.**
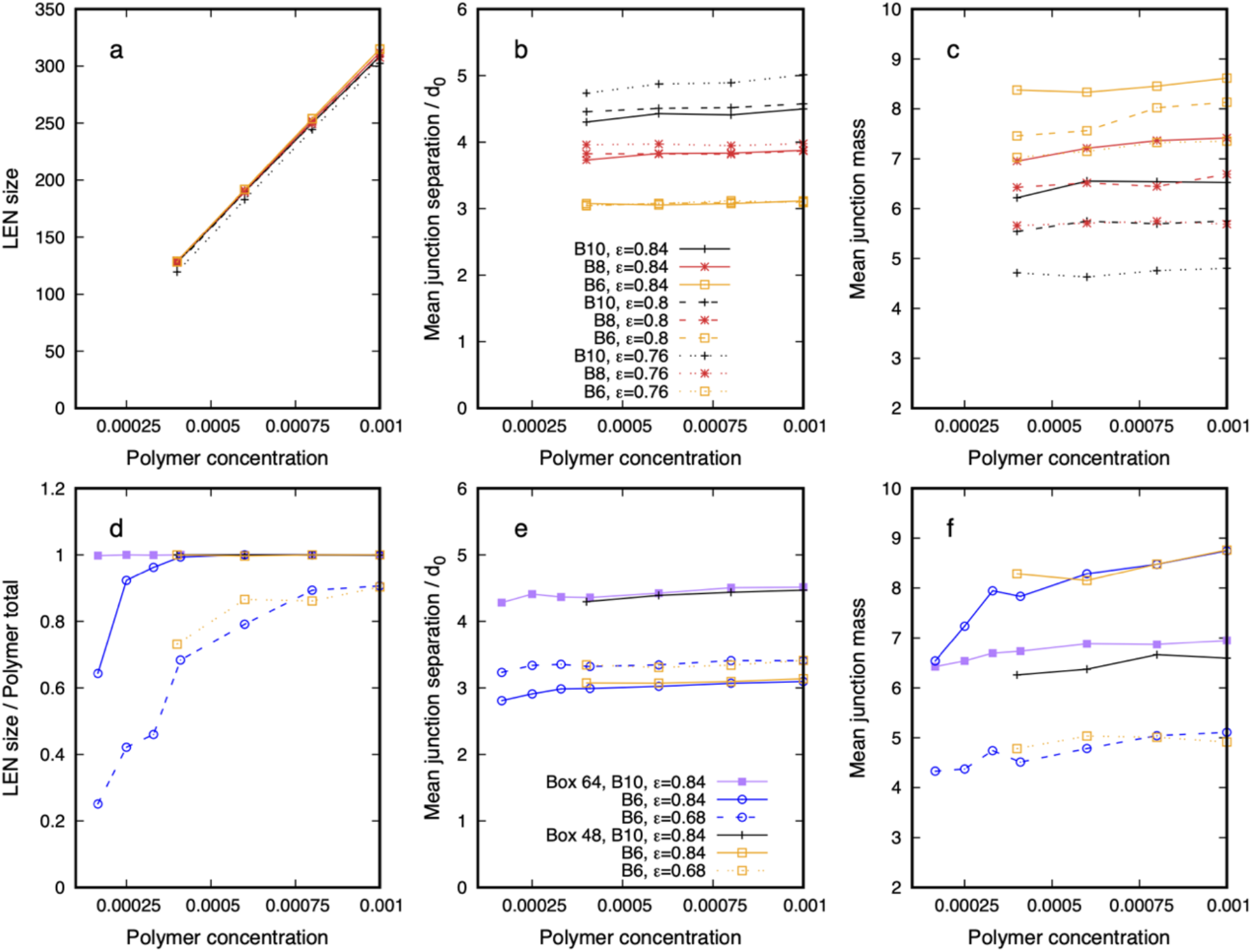
Quantitative properties of the dense phase of polymers 6Bm, m = 6, 8, 10, and three affinities, *ε* = 0.84, 0.8, 0.76, in a box (48 *d*_0_)^3^ (top row), and comparison of results for two affinities *ε* = 0.84, 0.68, in box sizes (48 *d*_0_)^3^ and (64 *d*_0_)^3^ (bottom row). (**a**) The Largest Equilibrium Network (LEN) size increases linearly with polymer concentration. (**b**) The junction separation increases with increasing binding site separation, but is largely independent of concentration and affinity. (**c**) The mean junction mass is independent of the polymer concentration, but shows a (small) systematic increase with increasing affinity and decreasing binding site separation. (**d**) The fraction of polymers in the LEN is independent of system size within the statistical accuracy of the simulations. This remains true even when ∼ 25% of the polymers remain in the dilute phase (dashed curves.) Both junction separation (**e**) and junction mass (**f**) are independent of system size and concentration. Statistical error bars are smaller than the symbol size. Each legend applies to all graphs in the row.

The top row of Figure 3 shows how properties of the dense phase of polymers 6Bm, m = 6, 8, 10, vary with the total concentration and binding site affinity in the standard simulation box (48 *d*_0_)^3^. The dense phase grows linearly with concentration (Panel 3a), a result that holds even for the lowest affinity and largest spacing (*ε* = 0.76, 10 beads, dotted black curve) for which more polymers remain in the dilute phase. The mean junction separation, shown in Panel 3b, increases when the binding sites are farther apart on the polymers, which is intuitively expected. But it is largely independent of the affinity, apart from a small divergence (less than the size of a single bead) for the 6B10 polymers, which suggests that the network is loosened by decreasing affinity (cp. bottom row of Fig. 2). This result implies that the porosity of a biomolecular condensate is insensitive to small changes in the strength of the interactions between the constituent IDPs but increases with increasing separation of their binding domains (cp. the top row of Fig. 2). Panel 3c shows that the mean junction mass increases systematically (albeit slowly) with increasing binding site affinity and decreasing separation. Combined with Panel 3b, we find that increasing the affinity while keeping their separation unchanged does not change the structure of the dense phase, but only packs more polymers between existing junctions.

We have verified that the dense phase is in equilibrium with the surrounding dilute phase by two methods. It is clear from panels 3b and 3c that the junction separation and mass are independent of the total polymer concentration in the range studied, which supports the droplets being in thermodynamic equilibrium. We also simulated systems in a larger box (64 *d*_0_)^3^ to confirm that the dense phase structure is independent of the simulation box size. Because the larger simulations are extremely time consuming, requiring 16 cpu-days per million time-steps, we have only compared two affinities (*ε* = 0.76, 0.84) and two separations (6, 10). Panel 3d shows the fraction of polymers in the droplet in both system sizes is independent of the box size even for weak affinities (dashed curves) for which 10 - 30% of the polymers remain in the dilute phase over this concentration range. Similarly, panels 3e and 3f show that the mean junction separation and mass are independent of the box size and total concentration for all binding site affinities and spacings studied. Note that the droplets in the larger simulation box have been simulated at three lower concentrations than those in the smaller box. These concentrations are chosen so that there are the same *number* of polymers in the larger box as for the first three concentrations in the smaller box. It is clear that while the structural properties of the droplets are the same (Panels 3e and 3f), Panel 3d shows that polymers migrate from the dense to the dilute phase when a well-formed droplet in the smaller box is simulated in the larger box, further supporting the equilibrium state of the system. Comparing the first data points of the dashed-blue/circle and yellow/square curves shows that while ∼70% of the polymers are in the dense phase in the smaller box, this falls to about 25% of the (same number of) polymers in the larger box. (Section 4 and Figure S10 of the Supplementary Material provides further supporting data.)

We find that model IDPs with multiple binding sites stretch on entering the dense phase, an effect that increases with increasing affinity. This counter-intuitive result may arise from its modified conformation ensemble in the two phases: while an IDP in dilute phase must double back to gain binding enthalpy, it can bind without bending to multiple junctions in the dense phase, thereby gaining binding enthalpy at the loss of a small amount of conformational entropy. We also find that although an IDP in the dense phase can, in principle, span the same pair of junctions multiple times (see Figures S7 and S8), this is actually quite rare. Only 16% of 6B6 polymers span the same junctions twice, and 1.6% three times (Table S1 in the Supplementary Material shows the fraction of single, double and triple spans of a junction). The model IDPs in the molecular dynamics simulations of Dignon *et al*., also stretch, showing this effect is robust against the simulation technique.(37)

The results of this section demonstrate that the model IDPs phase separate into a structured fluid in equilibrium with the surrounding dilute phase for a range of combinations of the location and affinity of their binding sites. This supports our comparing the properties of the ∼50 nm diameter simulated droplets with those of much larger experimental BCs. It also suggests that the cell can tolerate some variation in the strength of IDPs interactions without greatly modifying the stability and structure of their dense phase.

### C The dense phase has long-range spatial structure

The mean values of observables in Figure 3 do not reveal their spatial variation throughout the dense phase. The radial distribution function (RDF) of the binding sites gives more complete information. The RDF is the probability of finding two entities at a given separation in space. The RDF of the endcaps and internal binding sites for polymers of types 6B10 and 6B6 are shown in Figure 4 for *strong* and *weak* values of affinity (ε = 0.84, 0.76), where these terms are used qualitatively to describe the cases shown in the left and middle columns of Figure 2. The dense phase possesses spatial structure out to length scales 3 *d*_0_ − 8 *d*_0_ far beyond the monomer size *d*_0_. The peak near the origin counts binding sites present at the same junction, and is uninteresting. The first interesting peak in the RDF quantifies the separation of junctions that are spanned by polymers. Notably, it does not shift significantly when the binding site affinity is reduced, but increases as expected when their separation is increased. The second peak also moves to larger distances but the peaks are less pronounced for the weaker affinity than the stronger. The near coincidence of the peak heights for the solid and dashed curves of the same colour shows that the mean junction separation shown in Figure 3 is an average over junctions formed by endcaps and internal binding sites. The peak heights are proportional to the number of binding sites (4 internal and 2 endcaps), so dashed curves are always above the solid curves for the same affinity and spacing.

**Figure 4.**
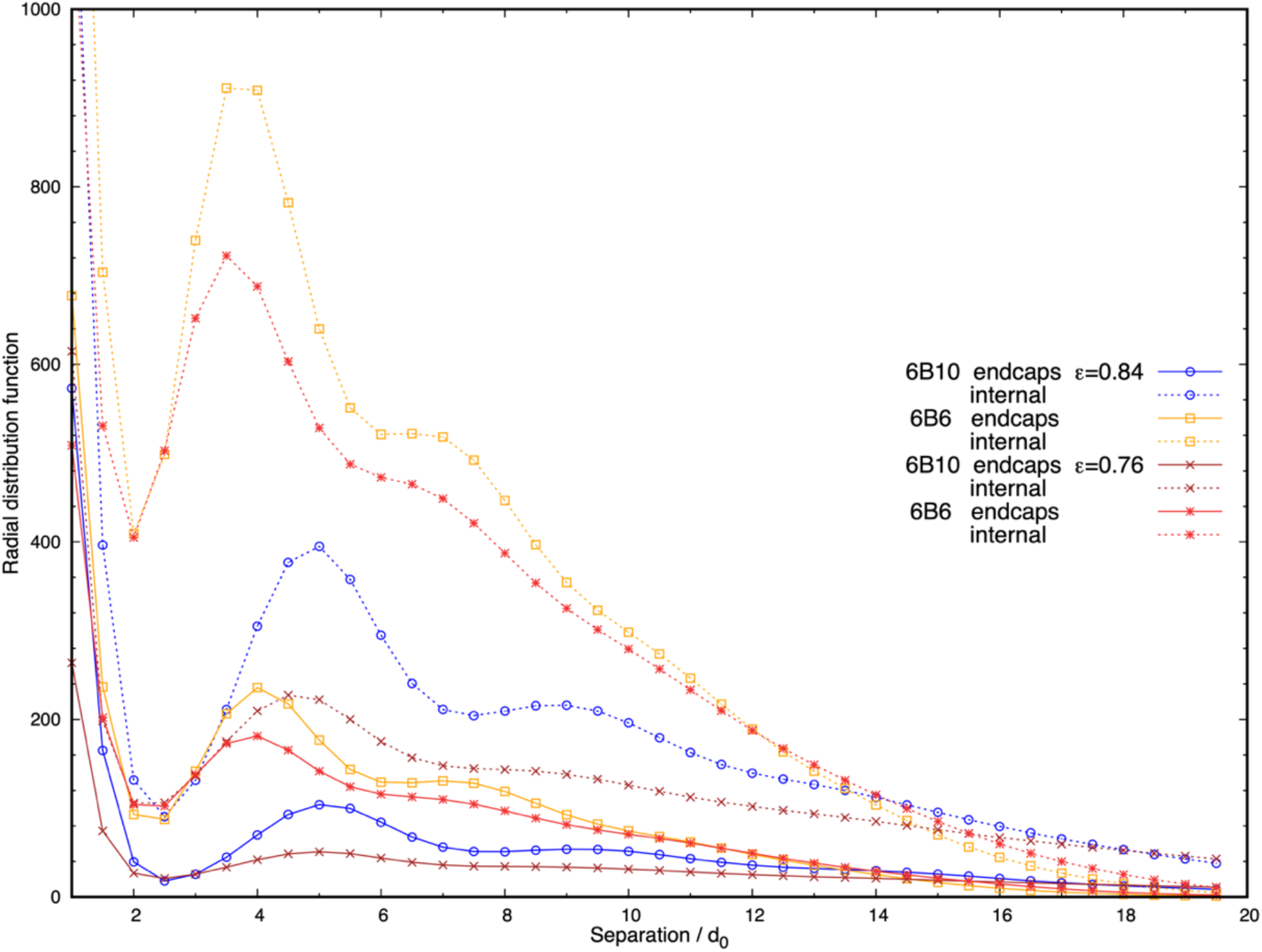
The radial distribution function of polymer binding sites in the dense phase reveals long-range organisation of junctions for two binding site affinities (ε = 0.84, 0.76) and separations (gap = 10, 6). Solid curves refer to endcaps and dashed curves to internal binding sites; colours distinguish the separation. The peak heights drop but are still distinguishable for 6B6 (yellow curve changes to red) and 6B10 (blue curve changes to brown) polymers on reducing the affinity from ε = 0.84 to 0.76.

### D Endcap binding sites are essential for phase separation

We have shown that the structure of the model biomolecular condensate responds to variations in the polymer binding site separation and affinity. This raises the question of what the consequences are of eliminating one or more of these sites. We performed independent simulations of 6B6 polymers with an affinity of *ε* = 0.84, over a range of concentrations, for the three cases in which either all sites are active, the two internal sites adjacent to the end-caps are deactivated, or the end-caps are deactivated. Turning off internal binding sites/endcaps in the simulations is accomplished by setting their conservative self-interactions/cross-interactions equal to their interaction with the water beads, which sets their affinity *ε* = 0. Note that the binding site beads are still present in the polymer, so its length is unaltered. Figure 5 shows typical droplets for these cases, in which the dense phase concentration (mean and standard deviation of 4 runs) is indicated below the snapshots. The first column shows that the phase separated droplet when all 6 binding sites are active has a concentration of ∼ 3 mM independent of the total polymer concentration. The droplet swells when two internal sites are turned off, and its concentration drops by a factor of 2-3 but it remains phase separated, as seen in the middle column. Note that if the dense phase is nearly spherical and crosses the periodic boundaries of the simulation box, this does not affect the calculation of its concentration. But the concentration of a non-spherical droplet is not easily measured, and the value for the bottom system in the middle column is included for completeness only. The final column shows that the dense phase disintegrates into hairy micelles or a box-spanning network with no sharp boundaries when the end-caps are disabled, even though the polymers still have 4 active binding sites.

**Figure 5.**
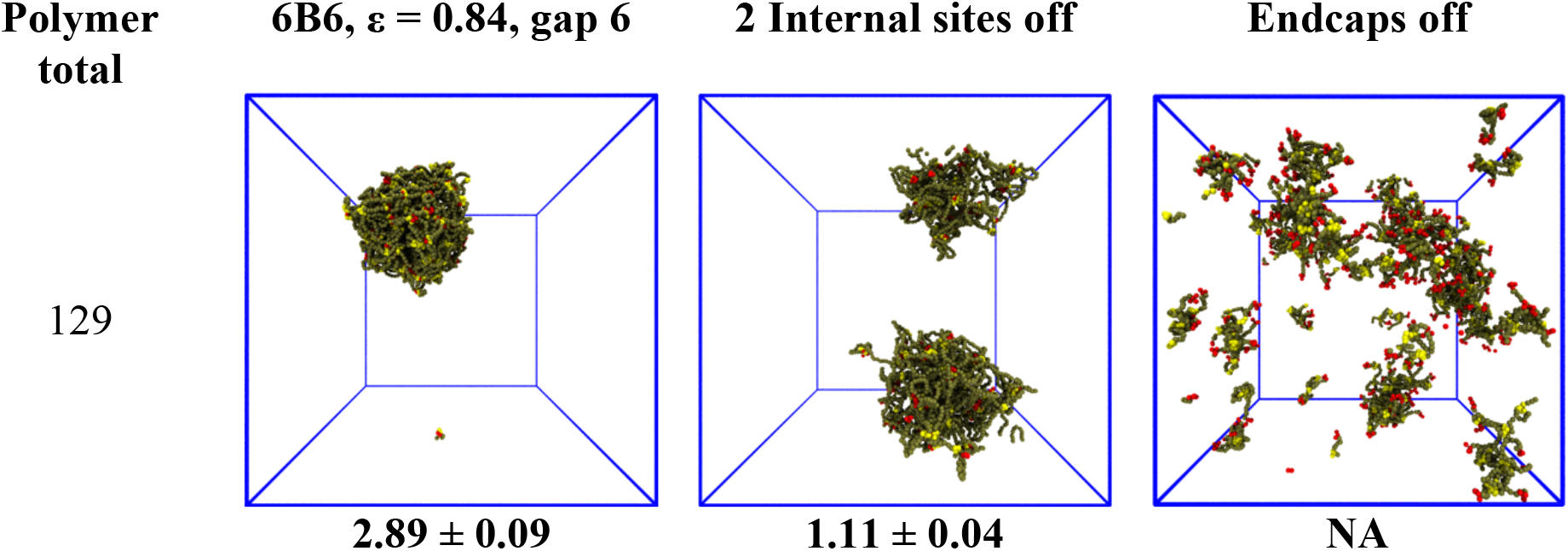
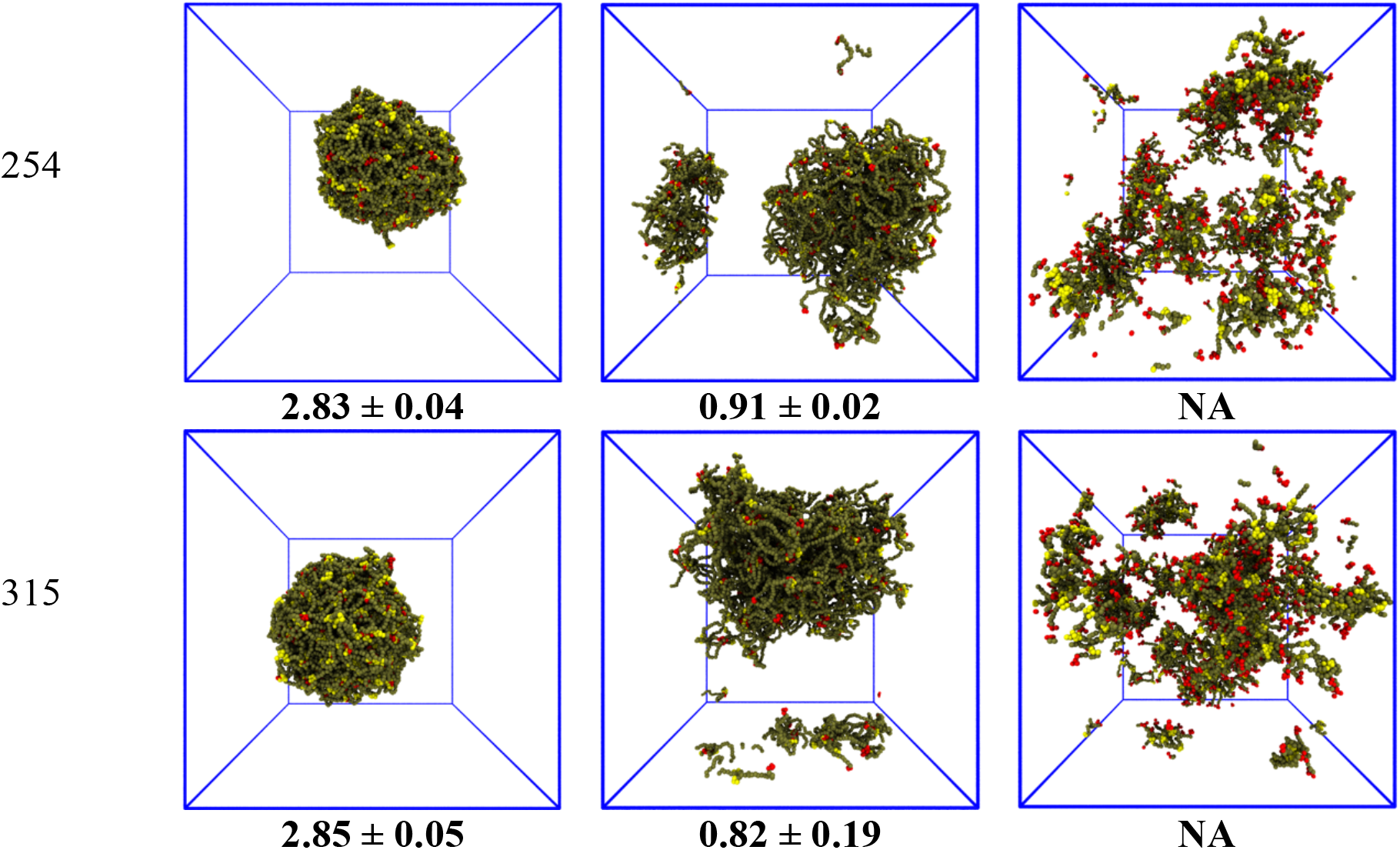
The dense phase responds differently to disabling internal binding sites or endcaps. Each row shows a droplet at a given total concentration (polymer total at left, numbers under snapshots are the dense phase concentration in mM averaged over four independent runs). In the first column, 6B6 polymers form stable droplets with an average concentration of ∼3 mM. The second column shows the systems with two internal binding sites turned off (i.e., polymer type 4B12-6-12). The droplet swells and the dense phase concentration decreases. The third column shows that without endcaps (i.e., polymer type 4I6) the droplet disperses. (NA = not applicable).

We quantify the change in the droplet structure when two internal sites are disabled in Figure 6 for polymers with three different separations (6B10, 6B8, 6B6) and fixed affinity *ε* = 0.84. Panel a shows that the network size changes little for the three cases, indicating that the polymers with fewer binding sites still phase separate into stable droplets. The mean junction separation, shown in Panel b, increases systematically by a factor that also increases with the binding site separation, reaching almost a factor of two when the binding sites are separated by 10 backbone beads. Concomitantly, the mean junction mass, in Panel c, decreases by an amount that is largest for the smallest binding site separation of 6 beads. We cannot measure dense phase properties when the end-caps are disabled because the remnant “networks” are too small to yield meaningful results.

**Figure 6.**
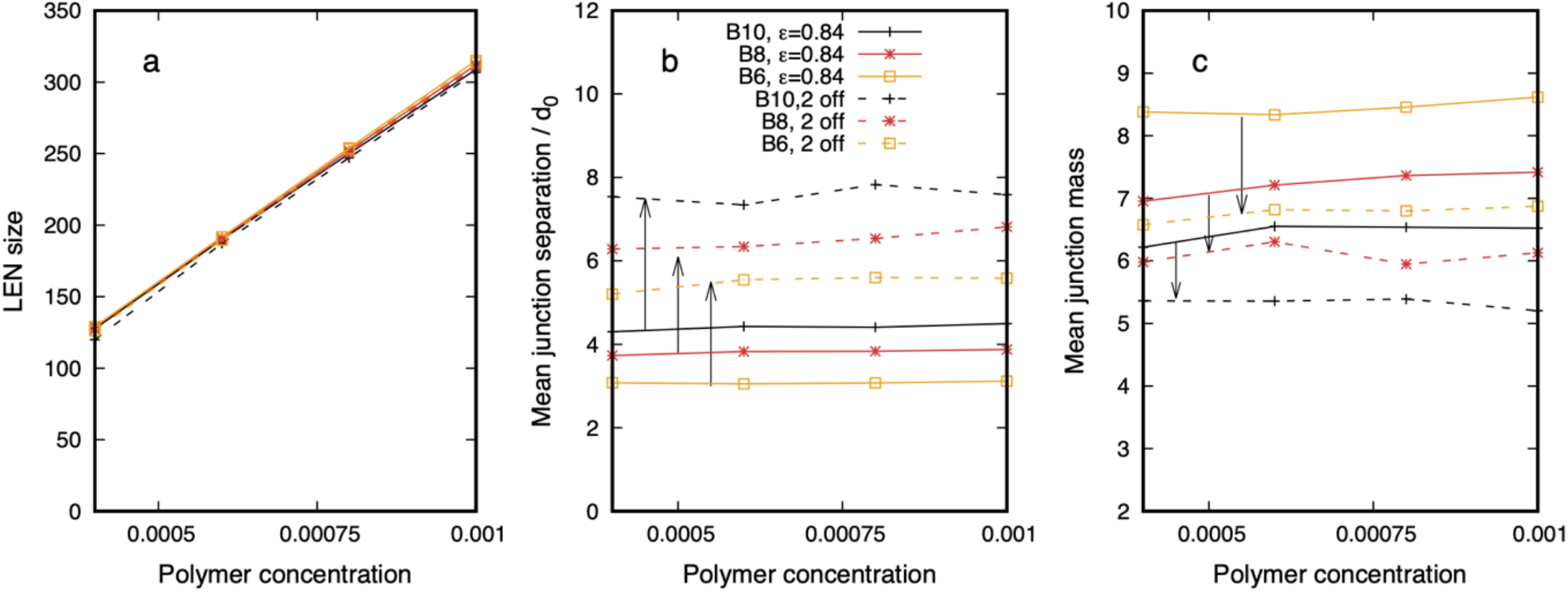
The structure of the dense phase of polymers with 6 binding sites and affinity ε = 0.84 (solid curves) changes when two inner sites are disabled (dashed curves). Arrows indicate the shift in the property. The binding site separation is colour coded – 6B10 (black), 6B8 (red), and 6B6 (yellow). (**a**) The dense phase contains (almost) all polymers for all separations and (internal) binding site states. The mean junction separation (**b**) increases, and the mean junction mass (**c**) decreases when the two sites are disabled. Snapshots corresponding to the solid and dashed yellow curves are seen in the left and centre columns in Figure 5. The legend applies to all.

Although results are shown for a single affinity (*ε* = 0.84), we expect from Figure 3 that disabling internal sites for other affinities would be similar unless the combination of binding site separation and affinity were such as to render the droplet already close to the phase boundary, e.g., 6B10 with *ε* = 0.76. Comparing Figures 3 and 6 shows that droplet’s characteristic length scale is more sensitive to the spacing of the binding sites than their affinity, while the junction mass is sensitive to both.

### E Clustering coefficient of the dense phase reveals a heterogeneous mass distribution

Although the model BC is an equilibrium thermodynamic phase, there is considerable variability in the spatial distribution of the mass. Figure 4 shows that the dense phase has a structure that extends far beyond the monomer size and is not smeared out over time in spite of its fluidity. The regulation of biochemical reactions has been identified as a key function of BCs, and the spatial organisation of scaffold proteins within a BC should be expected to influence the diffusion and reactivity of client molecules.(3, 91) Previous simulations of telechelic polymers has shown that the distribution of the number of polymers that bind at junctions throughout the dense phase is broad.(47) In this section we quantify the spatial heterogeneity of the dense phase using measures from graph theory.

The dense phase is mapped to a graph by identifying a *Node* with each junction where polymer binding sites meet, and connecting two nodes by an *Edge* when the corresponding junctions are spanned by at least one polymer. For the purpose of constructing the graph, an edge is assigned to two nodes if two adjacent binding sites on a polymer connect two junctions. Only polymers with at least two binding sites connecting them to the dense phase are included in the graph to avoid skewing its properties with polymers that dangle off into the dilute phase. The graph is recalculated each time the simulation is sampled to generate a time series of graphs that represent the droplet’s state from which equilibrium values of observables can be calculated.

All properties of the graph we use are calculated from its adjacency matrix *A*_*ij*_. This is the square matrix whose dimension is equal to the number of vertices in the graph, and whose elements *A*_*ij*_ = 1, if the vertices *i, j* are connected by an edge, and 0 if not. Diagonal elements are zero because we ignore self-loops, which represent polymers multiply bound to the same junction. The number of neighbours of a node *i*, referred to as its degree, is the sum *k*_*i*_ = Σ_*j*_ *A*_*ij*_ of the elements in row *i*.

From the matrix *A*_*ij*_ we construct the local Clustering Coefficient that measures the mutual connectivity of the nodes connected to a given node.(74, 75, 92) It is defined as the ratio of the number of linked pairwise neighbours of a node to the maximal possible number of linked pairwise neighbours averaged over all the nodes of the graph. The clustering coefficient *C*_*i*_ of node *i* is:

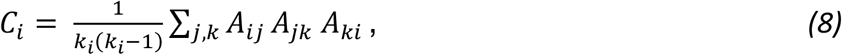

where *k*_*i*_ is its degree. The mean clustering coefficient (CC) is the average of *C*_*i*_ taken over all nodes in the graph. It measures the local connectivity of the nodes, taking its largest value of unity when all the neighbours of a node are also connected to each other (see Supplementary Material Section 1, Figures S1 – S6, for the CC of some simple graphs). The CC just defined is the unweighted clustering coefficient. We also define the weighted clustering coefficient (wCC),(75, 93) which is calculated using Eq. 8 with a modified adjacency matrix whose elements are *A*_*ij*_ = *n* if there are *n* edges between nodes *i, j*. The weighted clustering coefficient contains information about the number of polymers that connect two junctions, and provides a better measure of the mass distribution in the droplet than the unweighted CC that reflects only its connectivity.

Figure 7a shows that the CC of networks of polymers with multiple binding sites of affinity *ϵ* = 0.84 increases linearly with binding site number until it starts to saturate at six sites. The CC is 2-3 times larger than the value for a network of telechelic polymers (2B48) with a similar backbone length, and much greater than for a small-world random graph (see Section 2 in the Supplementary Material for the definition of small-world graphs). This indicates that multisite polymers form highly-connected junctions. The variance of the CC, indicated by its vertical extent, reinforces the evidence of the radial distribution function in Figure 4 that the local network density is heterogeneous. Because of the large parameter space of polymers with multiple binding sites, we have not calculated the CC for all combinations of their number and affinity. Section 5 of the Supplementary Material shows further data on the CC of the networks. For telechelic polymers, the CC increases approximately linearly with the binding site affinity (Figure S11). And for multisite polymers, the CC is independent of network size, and decreases with increasing binding site separation (Figures S12, S13). We find that that both the number, separation, and affinity of the binding sites influence the clustering coefficient.

**Figure 7.**
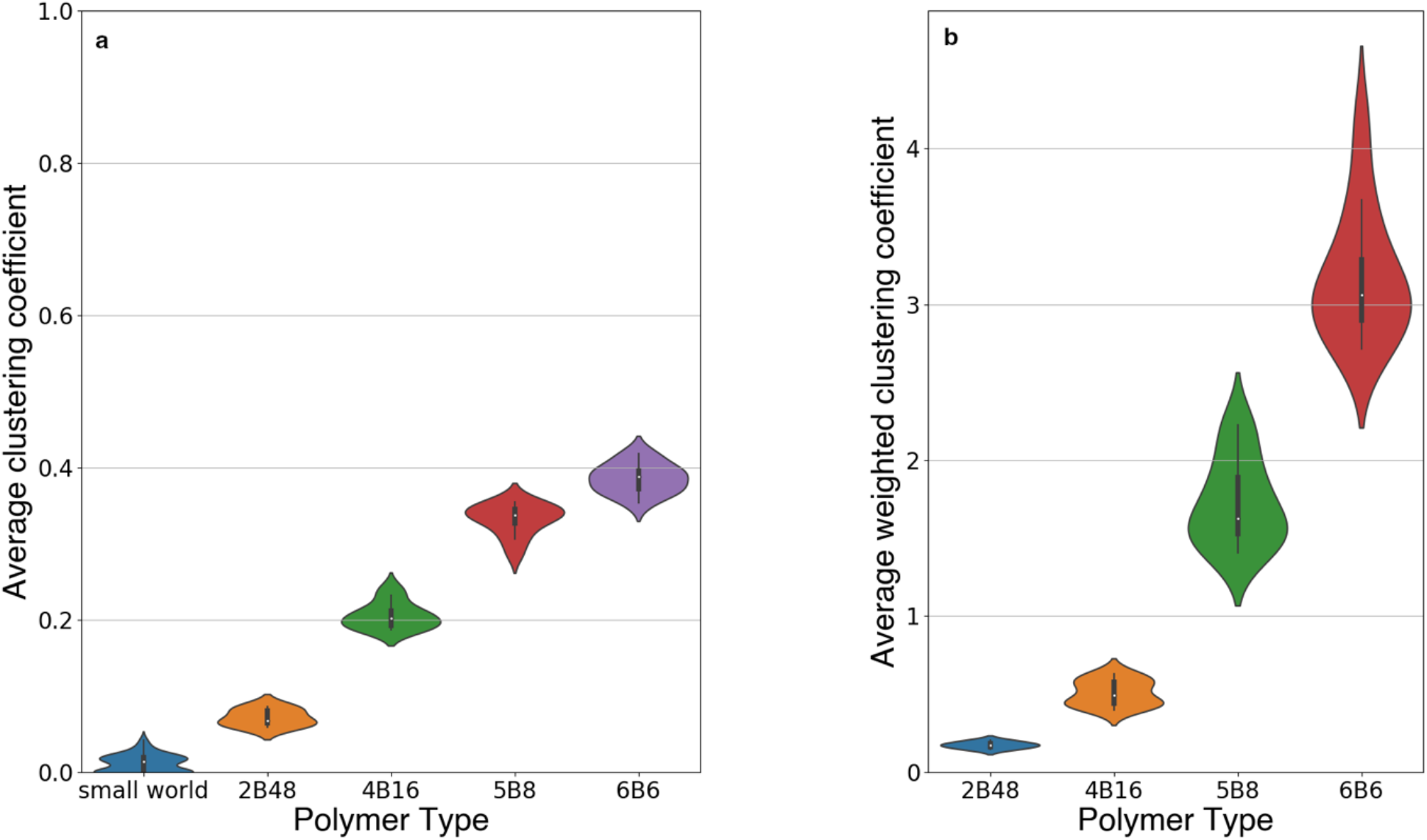
Unweighted (CC) and weighted (wCC) clustering coefficients for the dense phase of polymers with binding site affinity ε = 0.84, and 2 (2B48), 4 (4B16), 5 (5B8) and 6 (6B6) binding sites separated by decreasing lengths of linker beads to maintain the polymer length approximately constant. (**a**) The CC increases linearly with increasing numbers of binding sites. The left-most symbol is for a small-world random graph for comparison. (**b**) The wCC has a broader distribution than the unweighted CC, and its variance across the dense phase increases with more binding sites.

The wCC in Figure 7b shows that the average connectivity of junctions increases sharply with increasing binding sites per polymer and its variance increases particularly for polymers with 5 and 6 sites. Polymers with multiple binding sites form regions with a high local connection density as well as regions with a low density. These results are averaged over samples taken from simulations of 600,000 timesteps (after discarding 10^6^ steps) indicating that they persist in equilibrium. We point out here that the wCC is calculated for each node *i* in the graph and normalized by dividing by the quantity *k*_*i*_(*k*_*i*_ − 1), which is the maximum number of triangles the node could make, and then averaged over all nodes. The number of triangles a node actually makes is usually much smaller than this maximum (cp. Fig. 7a), which is why the ordinate of Fig. 7b is not large. But its vertical extent reflects the large variation in the number of polymers spanning the junctions. The wCC of the random graph is not shown because it has edges that span the whole graph, which renders it not comparable to locally-connected droplets. Although moving internal binding sites closer together along the IDPs systematically reduces the separation of the junctions in the BC (cp. Fig. 3b) and increases the mass distribution (cp. Fig. 3c), it remains highly heterogeneous.

We hypothesize that a cell may regulate the structure of a BC via post-translational modifications to scaffold proteins that occlude or expose interaction sites. To explore the prediction of our model in this case, we have compared the CC for the networks composed of 6B6 polymers with all six sites active and when two internal binding sites are disabled. These correspond to the networks shown in the middle column of Figure 5. Figure 8 shows that the CC for the network is largely independent of concentration for the four points A - D, but decreases when two internal binding sites are disabled (cp. “A” and “A two off”, etc.)

**Figure 8.**
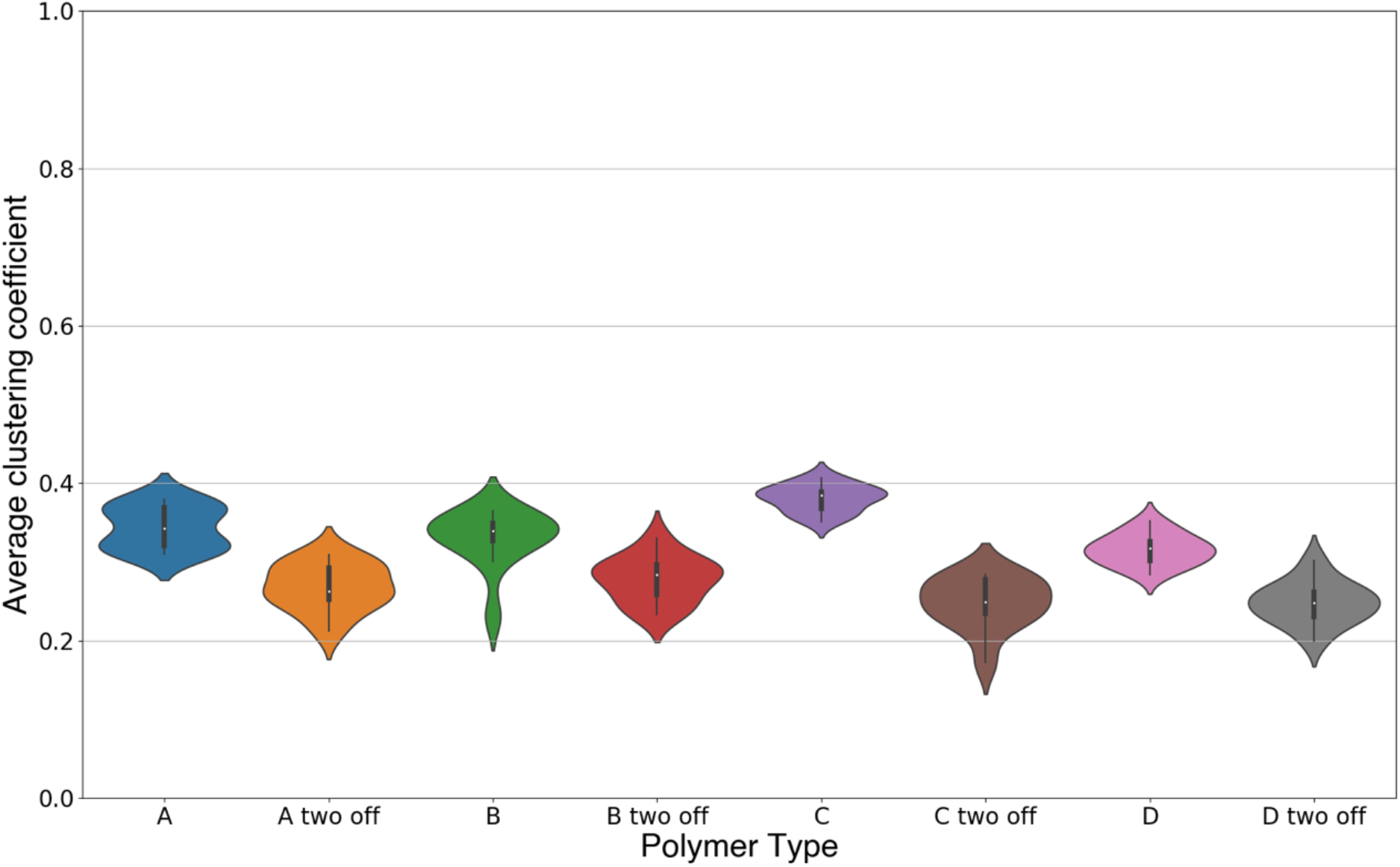
Comparison of the unweighted clustering coefficient for a network of 6B6 polymers with affinity ε = 0.84 and four concentrations (A = 129 polymers, B = 192, C = 254, D = 315) when all binding sites are active (A, etc.) and when two internal sites are disabled (A two off, etc.). The mean clustering coefficient is almost unchanged (∼ 4) for the four concentrations. Disabling two internal binding sites reduces the mean clustering coefficient (∼ 3) for all concentrations, but its variance remains similar, indicating that the network remains heterogeneous.

### F Binding site affinity and spacing influence dense phase fluidity

Many biomolecular condensates are in the fluid phase, and loss of fluidity has been proposed as a sign of pathology.(29) In this section, we demonstrate that the dense phase of the model IDPs is fluid, and quantify how its internal dynamics depends on the binding site number and affinity.

A common experimental demonstration of condensate fluidity is to follow the spontaneous fusion of two distinct droplets into a single, spherical droplet on contact.(32, 94) The same phenomenon is observed in our system. Figure 9 shows two droplets fusing, each of which is composed of 763 telechelic 2B8 polymers with an endcap affinity *ε* = 0.76. On encountering each other by diffusion, they merge and eventually form a single, spherical droplet. Supplementary Movie M6 shows the full evolution of the fusion event.

**Figure 9.**
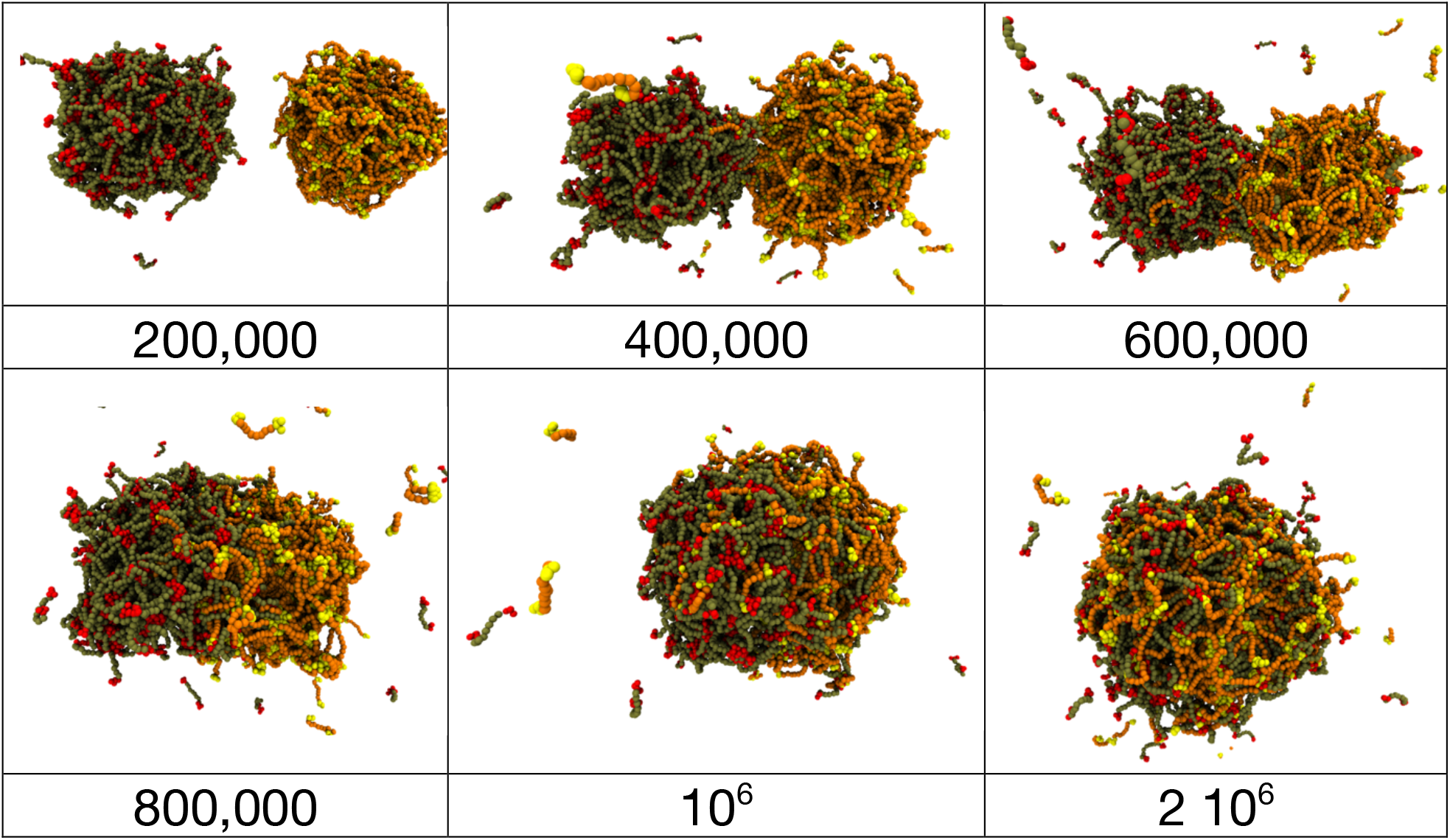
Fusion of droplets composed of 763 identical 2B8 polymers that are colored differently for clarity but are otherwise identical. After diffusing to contact, they merge and polymers diffuse throughout the combined droplet that evolves to a spherical shape indicating the presence of a non-zero surface tension. The simulation times at which the snapshots are taken are beneath the images. Note the increased interval between the last two snapshots.

The fusion of two droplets is a non-equilibrium process, but equilibrium droplets are not static. Polymers diffuse and exchange between the droplet and the dilute phase in an affinity-dependent manner. Polymer dynamics in the dense phase may intuitively be separated into the diffusion of the whole polymer, as measured by its (relatively slow) centre of mass (CM) motion, and (relatively fast) conformational fluctuations by which it can bind/unbind to/from other polymers. The latter processes occur even if the polymer’s CM is on average stationary. We introduce a new observable *f*(*τ*), which we call the network *fluidity*, that is defined on the droplet’s equivalent graph to display the conformational dynamics of the dense phase:

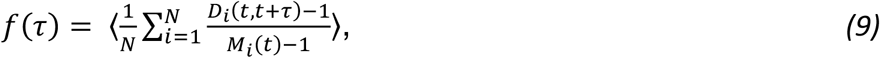

where *N* is the number of nodes in the graph, *M*_*i*_(*t*) is the number of polymer binding sites present on node *i* at time t, and *D*_*i*_(*t, t* + *τ*) is the number of descendent nodes at time *t* + *τ* that contain at least one binding site that was bound to node *i* at time *t*. Angle brackets indicate an average over all starting times t. The fluidity is normalized so that *f*(0) = 0 because each node is then its own descendent, *D*_*i*_(*t, t*) = 1. We note that this definition implies that if a polymer has more than one of its binding sites bound to a node and any of them move to a different node, they contribute to the fluidity measure. That is, the fluidity tracks the motion of the binding sites not the whole polymer. We also note that if a binding site leaves a junction and rebinds to the same one within the sampling time of the simulations (50,000 time steps) this is not reflected in the fluidity measure. It would appear if the sampling rate were increased. The function *f*(*τ*) measures the number of descendent nodes into which each node splits during a time interval *τ*, averaged over the whole graph and normalized so that if a node is its own descendent it contributes 0, and if it totally splits up it contributes 1. It is similar to the auto-correlation function of the number of binding sites present on a node with the important difference that it tracks not only the *number* of binding sites per node but also their *identity*: if a binding site leaves a node and a different one joins, the fluidity measure reflects this even though the *number* of binding sites on the node has not changed. We introduce this definition because binding sites fluctuating between nodes represents a motion that will likely impact the behaviour of other molecules within it, e.g., by permitting the diffusion of objects larger than the mean junction separation because the network can rearrange around them.

The fluidity of the dense phase of telechelic 2B16 polymers is shown in Figure 10a for a range of binding site affinities. These polymers form box-spanning networks for the lowest two affinities *ε* = 0.76, 0.74, and phase separated droplets for higher values.(47) The fluidity function for weak affinity increases steeply over short times revealing that the polymers easily rearrange their binding configuration as they diffuse within the dense phase. As the affinity increases, the droplet dynamics slows down. The low-affinity networks rearrange their structure after a few hundred thousand timesteps, whereas the high affinity polymers require millions of steps, which means these droplets evolve slowly on the time scale of the simulations. Table 2 shows the timescale for the dynamic rearrangement of the networks shown in Figure 10.

**Figure 10.**
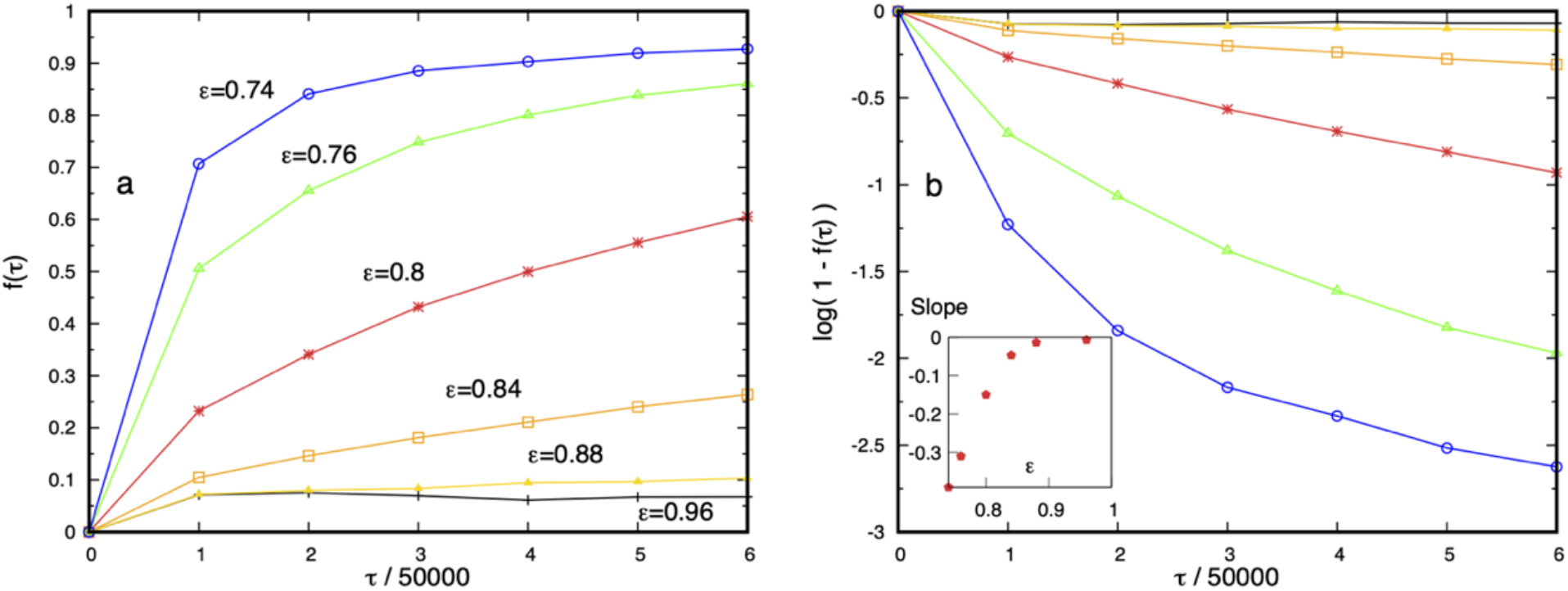
**(a)** The fluidity function *f*(*τ*) quantifies the ease of polymer binding sites fluctuating between junctions in the dense phase. For 2B16 telechelic polymers, it increases strongly with decreasing binding site affinity. The lowest two affinities (*ε* < 0.8) yield box-spanning networks while higher affinities form phase-separated droplets. The low fluidity for the highest affinities indicates that polymers fluctuate slowly on the time-scale of the simulations. (**b**) Semi-logarithmic plot of the fluidity of the 2B16 polymers showing that their dynamics are dominated by a single relaxation timescale for high affinities, while the lowest two affinities exhibit more than one timescale. Inset shows the slopes obtained from linear regression. The two affinities below 0.8 are included for completeness although the curves are not linear.

**Table 2.** The relaxation time of the fluidity function *f*(*τ*) for droplets of telechelic polymers with different endcap affinities. It is extracted from the slope of the curves in Figure 10b assuming single exponential decay. Only values for which the curves are initially linear in the figure should be considered accurate, but the lower affinity values show that the box-spanning networks evolve rapidly.

| Endcap affinity $\epsilon$ | Relaxation time / $10^6$ steps |
| --- | --- |
| 0.96 | 7.2 |
| 0.88 | 3.6 |
| 0.84 | 1.05 |
| 0.8 | 0.34 |
| 0.76 | 0.16 |
| 0.74 | 0.13 |

The curves for affinities above *ε* = 0.8 are well fitted by a single exponential in Figure 10b, indicating that their dynamics is dominated by a characteristic timescale. The fluidity of droplets composed of polymers with weaker affinity cannot be described by a single timescale implying more complex dynamics. The inset shows the slope of the fluidity versus time curve obtained by fitting the data in the semi-logarithmic plot of Figure 10b to a single exponential decay and extracting the slope from linear regression of all data points. The lowest two affinities are included for completeness although the curves are clearly not linear. We hypothesize that the dynamics of high-affinity networks is dominated by the binding-unbinding times of the polymers and not their spatial diffusion, whereas at least two time-scales contribute for lower affinities. Recent, molecular dynamics simulations and theory also predict complex dynamics inside BCs that depends on the stoichiometry between two types of multisite associative polymers.(95)

Adding more binding sites might be expected to reduce the dense phase fluidity, but this is not observed. Figure 11 shows that droplets with multiple binding sites are more fluid than telechelic polymers (Figure 10) with the same affinity *ε* = 0.84, and exhibit fluctuations on shorter timescales. Figure 11a shows that the dynamics, as measured by the fluidity *f*(*τ*), slows down with increasing affinity as intuitively expected, but that the fast motion of the polymers is reduced with increasing binding site separation for stronger affinities (dashed 6B6 curves are above solid 6B10 curves). We attribute this effect to the increasing difficulty of the binding sites on a fluctuating polymer to move to a nearby junction as their separation along the polymer backbone increases (cp. junction separation in Figure 3). Figure 11b shows that the fluidity of droplets containing multi-binding site polymers cannot be described by single exponential decay for any of the affinities studied. Their internal dynamics therefore possesses at least two distinct time-scales. The departure of the curves for 6B10 polymers with the lowest affinity and concentration reflects the small size and instability of the dense phase.

**Figure 11.**
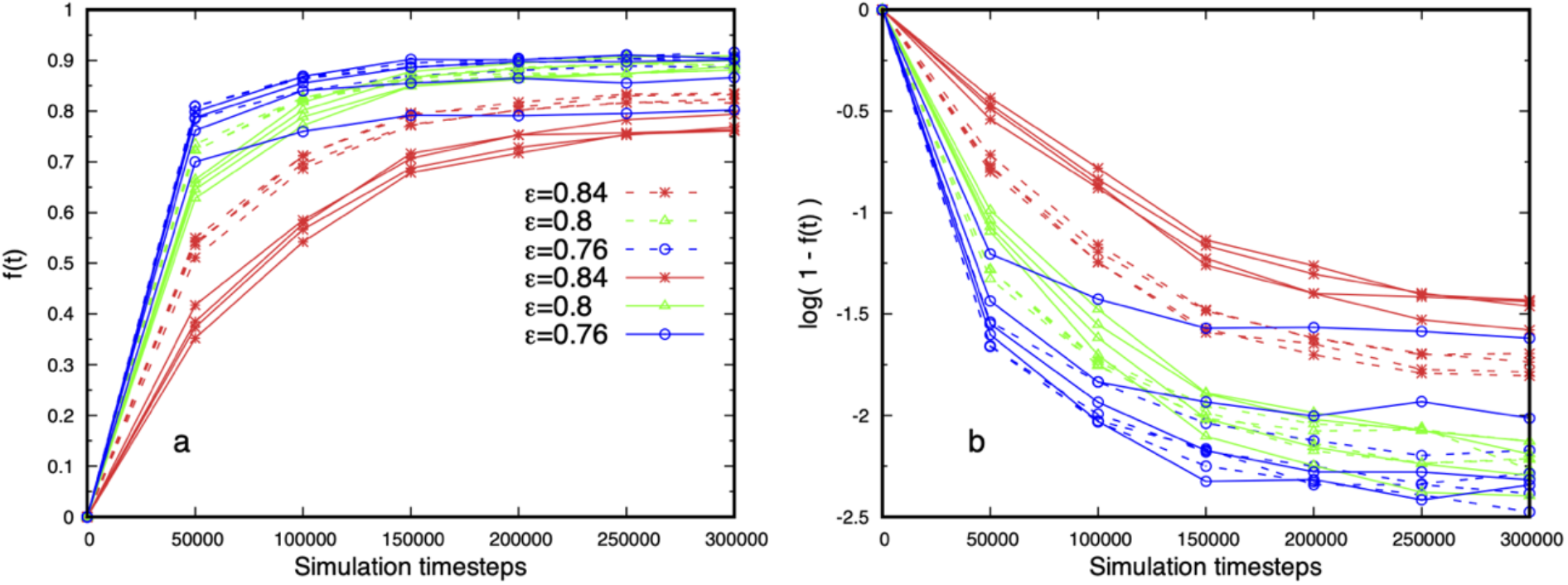
(**a**) Fluidity of droplets formed of 6B10 (solid) and 6B6 (dashed) polymers with 6 binding sites of three affinities. Repeated curves of the same colour and type are from simulations at four concentrations. The fluidity increases with decreasing affinity, and is independent of concentration except for the lowest affinity with the larger binding site separation (solid blue curve), indicating that the droplets are in equilibrium. The 6B10 polymers are less fluid than the 6B6 for higher affinities. (**b**) Log-linear plot of the data showing that the fluidity cannot be described as an exponential decay with a single time-scale for the affinities shown.

Finally, the independence of the fluidity on the total polymer concentration, and therefore on the droplet size, seen in Figure 11 further supports the claim that the droplets are in equilibrium. Snapshots of droplets corresponding to Figure 11 for the affinities *ε* = 0.8, 0.76 are shown in the left and middle images in Figure 2, and quantitative measures of their structure in Figures 3 and 4.

## Discussion

The Flory-Huggins (FH) theory of polymer phase separation(89) has been widely used to construct and interpret simulation studies of IDP phase behaviour.(37, 38, 45, 61, 96) This theory assigns all monomers in a polymer an energetic interaction with the solvent that is quantified by a parameter χ (for heteropolymers this is an effective parameter). When χ is negative, the entropy of mixing combines with the favourable solvation energy to keep the single-phase mixture stable. This corresponds to the *good solvent* limit. As the parameter χ is increased from a negative value, there is a point at which the increasingly repulsive polymer-solvent interactions precisely balance the entropy of mixing, and the polymers fluctuate as ideal chains. This defines the *theta point* of the mixture. As χ becomes more positive, an initially homogeneous mixture of polymer and solvent spontaneously separates into a polymer-rich phase and a polymer-poor phase, yielding the *poor solvent* limit. Phase separation only occurs in FH theory when all (or a large fraction) of the monomers in the polymers possess a sufficiently solvophobic interaction (large positive χ parameter).(38, 45)

We have here explored a quite different model of IDPs as polymers in good solvent conditions (*i*.*e*., with a zero or negative χ parameter) to understand IDPs that do not contain a majority of hydrophobic residues (*e*.*g*., FUS-LC is almost uncharged and has only ∼ 25% hydrophobic residues). The phase separation and dense phase properties we observe cannot be understood within FH theory because the intermolecular attractions are too weak to collapse the polymers in the good solvent. Although inspired by biological IDPs, our model is applicable to the phase separation of uncharged, hydrophilic polymers possessing punctate sticky sites, and its extension to multicomponent systems is obvious.(97-100)

In the good solvent regime, a single polymer exhibits Flory scaling in which its end-to-end length scales with the number of monomers as *L*_*ee*_ = *aN*^*ν*^, where ν = 0.5 for an ideal chain and 0.6 for a self-avoiding chain, and the prefactor a is related to the monomer size.(89) Tomasso *et al*.(87) present data for a wide range of IDPs showing that their hydrodynamic radius in dilute solution indeed scales close to an ideal chain, with the exponent ν = 0.509. These IDPs have the conformational fluctuations typical of polymers in a theta or good solvent, and cannot phase separate without additional attractive forces. A simple example of such forces is provided by telechelic polymers, in which the endcaps have an attractive self-interaction while both endcaps and backbone are solvophilic, i.e., their χ parameter is negative.(97) Theory(98, 99) and computer simulations(47, 101-104) show that telechelic polymers phase separate into a reversible physical gel.(89)

We have shown that model IDPs with punctate attractive binding sites phase separate into a dense phase in equilibrium with a dilute phase in the absence of any solvophobic repulsion. The dense phase has a concentration in the range 2-12 mM, and an aqueous volume fraction in the range 60-70 % when the simulated IDPs are mapped to the FUS low complexity domain, in good agreement with experimental results.(33) The dense phase is a dynamic network, in which IDPs binding sites transiently assemble at junctions (or nodes), and resembles associative polymer systems.(97, 105, 106) The spatial structure (or porosity) of the dense phase depends strongly on the IDP’s binding site separation but less so on their affinity (Figure 3), and has a characteristic length scale much larger than the length scale of the monomers (Figure 4). This length scale is selected by the separation between junctions (where binding sites meet), and is generally reduced on increasing the number of binding sites, but it is not simply related to the density, as opposed to molecular fluids. The presence of multiple binding sites extends the range of affinity and IDP backbone length for which phase separation is observed, a robustness that may be important for cellular control of BCs composed of IDPs of variable size and imprecise composition. We may contrast the dense phase structure we observe with the results of Statt *et al*. who used coarse-grained Molecular dynamics simulations to map the temperature/density phase diagram of a model IDP.(38) In their simulations, IDPs were modelled as amphiphilic polymers composed of 20 hydrophobic and hydrophilic beads whose sequence and number were varied, but maintaining at least 60% hydrophobic beads. Statt *et al*. observed a variety of dense phase morphologies, ranging from a mixed fluid, to micellar, wormlike micellar, and lamellar phases as the temperature was changed. However, the radial distribution function of their dense phase showed no structure beyond 2-3 bead diameters for all sequences studied indicating an homogeneous dense phase, while we observe structure out to 10 bead diameters (Figure 4).

We next explored how the phase separation is affected when a fraction of the binding sites are disabled (Figure 5). A clear difference is found when sites distant from the polymer endcaps are disabled (dense phase swells and its concentration drops) compared to sites at the endcaps (dense phase dissolves.) Extrapolating our results to experimental IDPs, we predict that disabling interaction domains near their termini, by mutation or posttranslational modification, will destabilise the dense phase. But disabling domains within the IDPs is likely only to modulate the dense phase concentration. It is worth noting here that PTMs often cluster at the termini of IDPs. For example, FUS contains multiple PTM sites in its N-terminal prion-like domain (PLD) that are phosphorylated under stress conditions and reduce its tendency to undergo phase separation.(107) Phosphomimetic alterations of the PLD also influence the transition of FUS granules into pathological solid phases, while leaving the dynamics in the fluid stress granules unaltered.(108) The conformational flexibility of IDPs makes these terminal PTM sites easily accessible to diffusing species in the cytoplasm suggesting that regulation by diffusing kinases and phosphatases may be important for the function of FUS. Our results suggest that the reaction and diffusion of client proteins inside biomolecular condensates could by modulated by turning on or off appropriate interaction domains on their scaffold proteins, thereby modifying its porosity. It is instructive to compare these results to recent Monte Carlo simulations of Rana *et al*. whose model has a similar level of coarse-graining.(62) An IDP is represented in their work as a sequence of (hydrophobic) sticker beads interspersed with (hydrophilic) spacer beads, in which at least 40% of the polymer sequence must be hydrophobic in order to observe phase separation. Our model IDPs are wholly hydrophilic and attract each other only at punctate binding sites. Rana *et al*. find that the phase behaviour of their model is sensitive to the identity of the terminal beads, as we do, but their system changes from phase separated to aggregated clusters under the change of identity whereas we predict that the dense phase should dissolve.

The simulated BCs are fluid yet maintain an heterogeneous mass distribution over time. By mapping the junctions to nodes of a graph, we used the clustering coefficient (CC) to measure the connectivity between nodes in the network and the weighted CC (wCC) for their mass distribution. Both the CC and the variance of the wCC (measured throughout the BC) increase strongly with increasing binding site number, indicating a heterogeneous, although still fluid, mass distribution (Figure 7). We have constructed a novel measure (Eq. 9) to quantify the dynamics of polymer fluctuations within the dense phase. It reflects the reversible binding/unbinding of the IDPs, and shows that they exhibit multi-exponential decay in which both fast and slow relaxation processes are important (Figure 11). The relaxation dynamics we find is almost independent of IDP concentration and is not accompanied by stiffening of the network structure in the presence of more (weak) internal binding sites.

Fluid FUS-LC droplets have been observed to undergo an irreversible transition to a rigid fibrous state over several hours *in vitro*.(32, 109), and passive rheology experiments show that BCs undergo ageing and exhibit glassy behaviour.(110) However, condensates with material properties that range from liquid to solid also have physiological roles in healthy cells.(111) Alpha synuclein (aSyn) fibrillisation is preceded by phase separation into dense fluid droplets, but the individual aSyn molecules are more rigid than in the dilute phase.(112) The complexity of the electrostatic and hydrophobic domains in aSyn suggest that more complex forces are important in this case. On the other hand, small molecule drugs that harden biomolecular condensates can prevent RNA virus replication and provide novel therapeutic possibilities.(113) The relations between the dense phase dynamics and IDP binding site location/affinity that we predict may be relevant for cells that must robustly regulate BCs of variable size and composition. But they imply that increasing the affinity of multiple interaction domains will constrain proteins’ conformational ensembles, thereby accelerating their transition into pathological amyloid-forming conformations.

## Conclusions

The formation of biomolecular condensates via liquid-liquid phase separation (LLPS) of proteins has become a new paradigm within which to explain a wide range of phenomena in cellular physiology and pathology. Fluid macromolecular assemblies provide a segregated environment for cellular processes in the absence of membranes. But little solid information about their structure and dynamics is yet available, except for the primary sequence of the participating IDPs and the fact that under appropriate conditions they phase separate. Nevertheless, we can conclude based on the data that the widely-invoked explanation of LLPS based on the Flory-Huggins (FH) theory of polymer solutions fails for some IDPs, since they display *good solvent* properties in cytosol buffer conditions. We highlight concrete suggestions for experimental verification of our predictions, as well as the generalization of our analysis to other fields.

Although the model IDP binding sites do not represent specific residue-residue interactions, our finding of the special role of the end-caps is especially interesting in the light of experimental observations that post-translational modification sites are often clustered along the disordered termini of IDPs.(107) Our first prediction is that location and separation of interaction domains on phase-separating IDPs are key parameters for BC structure, but their precise affinity is not, except that the end-cap affinity is indispensable, *i*.*e*., phase separation can take place with binding end-caps in the absence of internal binding sites, while the opposite does not hold. This could be experimentally tested by generating mutant IDPs, such as FUS, with more inert residues at the termini and observing their phase behaviour. A second prediction is that varying the spacing between attractive domains in IDPs modulates the dense phase structure, and Small Angle Neutron Scattering (SANS) in conjunction with specific deuteration and contrast variation techniques is a promising tool to determine if the predicted scaling holds for experimental BCs.

Many IDPs possess multiple PTMs that can be modified by kinase/phosphatase species diffusing through the dense phase. A cell may control LLPS using post-translational modification to adjust the separation and/or affinity of attractive residues.(114) Steps towards computationally predicting how PTMs affect BC structure and dynamics have already been taken.(115) Given that the diffusion of enzymes would respond to the changed environment, this provides a level of feedback to the cell that could be important for regulating biochemical reactions within BCs. We have shown that the dense phase dynamics slows down on increasing the attraction between IDPs, and our third plea for experimental testing is to hasten measurements of dynamic properties of engineered BC phases. Fluorescence recovery after photobleaching could be used to test our prediction of the dependence of BC fluidity on binding domain strength.(116) Single-particle tracking experiments that follow the diffusion of variously-sized fluorescent probes, or quantum dots, within the BC would probe the predicted porous nature of BCs.

Although time-varying graphs are common in many fields, there are not yet many good tools for studying them. The graph fluidity measure we introduce could be applied to bioinformatics networks, such as genome association in dynamic complex traits,(117) metabolic networks,(71) and be combined with molecular dynamics simulations of large flexible proteins to explore their allosteric properties.(72, 73) It may also be useful for studying the evolution of social networks,(118, 119) the emergence of a power-law distribution for the node degree and the process of rumour spreading,(120, 121) as well as transport and mobility networks, which typically involve shortest-path and trajectory prediction on time-varying graphs.(122, 123)

In conclusion, the predicted relations between the model IDPs and the structure of their condensed phase will, if confirmed in experiments, help the design of synthetic BCs with tunable properties.(124-126) Combined with the ability to modify the residue sequence of IDPs,(60) and online tools to predict the effects of point mutations on local disorder and hydropathy of IDPs,(127) this will undoubtedly reveal more fascinating mysteries in cellular use of BCs,(128) and further our understanding of their pathological transitions in disease.(129)

## Author contributions

**J. C. Shillcock**, Conceptualization, Methodology, Software, Validation, Formal analysis, Data curation, Visualization, Writing – original draft, Writing – review and editing;

**C. Lagisquet**, Software, Validation, Data curation, Visualization, Writing - review and editing;

**J. Alexandre**, Software, Validation, Formal analysis, Writing - review and editing.

**L. Vuillon**, Software, Validation, Formal analysis, Resources, Writing - review and editing, Funding acquisition, Project administration.

**J. H. Ipsen**, Conceptualization, Methodology, Resources, Writing – review and editing, Project administration.

## Acknowledgements

This study was supported by funding to the Blue Brain Project, a research centre of the École polytechnique fédérale de Lausanne (EPFL), from the Swiss government’s ETH Board of the Swiss Federal Institutes of Technology, and the 80PRIME MITI CNRS programme. The authors gratefully acknowledge computer time provided by the Blue Brain Project and Swiss National Supercomputing Centre and the Abacus 2.0 Supercomputing cluster at the University of Southern Denmark. The Dissipative Particle Dynamics simulation source code used in this work is available on GitHub.(76) Custom python code written for the data analysis includes algorithms from the open-source libraries scikit-learn (https://scikit-learn.org/stable/) and networkx (https://networkx.org). The graph analysis code is available on GitHub (https://github.com/clagis/droplets). The authors express their gratitude to M. Brochut and E. Chénais who wrote the analysis code for Ref. 47, which was also used here, and to W. Pezeshkian for scripts to generate VMD format files. Snapshots and movies of simulated networks were produced using the open-source VMD software from the University of Illinois Urbana Champaign (http://www.ks.uiuc.edu/Research/vmd/).(130)

## Supporting Information

The Supporting Information is available free of charge *via* the internet, and includes the following:

> Technical details of the graph theoretic measures used, supporting results for condensed networks of polymers with a range of backbone lengths and end-cap affinities; data showing that the dense phase concentration is independent of the system size, and that it is highly solvated. Movies of the networks under different conditions are also provided.
>
> Movie M1 Equilibrium fluctuations of 315 polymers of type 6B6 with *ε* = 0.84.
>
> Movie M2 Equilibrium fluctuations of 309 polymers of type 6B10 with *ε* = 0.84.
>
> Movie M3 Equilibrium fluctuations of 315 polymers of type 6B6 with *ε* = 0.74.
>
> Movie M4 Equilibrium fluctuations of 190 polymers of type 6B10 with *ε* = 0.74.
>
> Movie M5 Equilibrium fluctuations of 254 polymers of type 6B6 with *ε* = 0.68.
>
> Movie M6 Fusion of droplets each containing 763 polymers of type 2B8 with *ε* = 0.76.

## References

1. Zhang JZ, Mehta S, Zhang J. Liquid–liquid phase separation: a principal organizer of the cell’s biochemical activity architecture. Trends in Pharmacological Sciences. 2021;42(10):845–56.

2. Holehouse AS, Pappu RV. Functional Implications of Intracellular Phase Transitions. Biochemistry. 2018;57:2415–23.

3. Lyon AS, Peeples WB, Rosen MK. A framework for understanding the functions of biomolecular condensates across scales. Nature Reviews Molecular Cell Biology. 2021;22(3):215–35.

4. Banani SF, Lee HO, Hyman AA, Rosen MK. Biomolecular Condensates: Organizers of Cellular Biochemistry. Nature Rev Mol Cell Biol. 2017;18:285–98.

5. Stroberg W, Schnell S. Do Cellular Condensates Accelerate Biochemical Reactions? Lessons from Microdroplet Chemistry. Biophysical Journal. 2018;115:3–8.

6. Seif E, Kang JJ, Sasseville C, Senkovich O, Kaltashov A, Boulier EL, et al. Phase separation by the polyhomeotic sterile alpha motif compartmentalizes Polycomb Group proteins and enhances their activity. Nature Communications. 2020;11:5609.

7. Zhang Y, Narlikar GJ, Kutateladze TG. Enzymatic Reactions inside Biological Condensates. Journal of Molecular Biology. 2021;433(12):166624.

8. Saha B, Chatterjee A, Reja A, Das D. Condensates of short peptides and ATP for the temporal regulation of cytochrome c activity. ChemComm. 2019;55:14194–7.

9. Dzuricky M, Rogers BA, Shahid A, Cremer PS, Chilkoti A. De novo engineering of intracellular condensates using artificial disordered proteins. Nature Chemistry. 2020;12(9):814–25.

10. Heidenreich M, Georgeson JM, Locatelli E, Rovigatti L, Nandi SK, Steinberg A, et al. Designer protein assemblies with tunable phase diagrams in living cells. Nature Chemical Biology. 2020;16:939–45.

11. Deviri D, Safran SA. Physical theory of biological noise buffering by multicomponent phase separation. Proceedings of the National Academy of Sciences. 2021;118(25):e2100099118.

12. Klosin A, Oltsch F, Harmon T, Honigmann A, Jülicher F, Hyman AA, et al. Phase separation provides a mechanism to reduce noise in cells. Science. 2020;367(6476):464–8.

13. de Curtis I. Biomolecular Condensates at the Front: Cell Migration Meets Phase Separation. Trends in Cell Biology. 2021;31(3):145–8.

14. Milovanovic D, Wu Y, Bian X, Camilli PD. A Liquid Phase of Synapsin and Lipid Vesicles. Science. 2018;361:604–7.

15. Hoffmann C, Sansevrino R, Morabito G, Logan C, Vabulas RM, Ulusoy A, et al. Synapsin Condensates Recruit alpha-Synuclein. Journal of Molecular Biology. 2021;433(12):166961.

16. Zeng M, Chen X, Guan D, Xu J, Wu H, Tong P, et al. Reconstituted Postsynaptic Density as a Molecular Patform for Understanding Synapse Formation and Plasticity. Cell. 2018;174:1172–87.

17. Wu X, Cai Q, Feng Z, Zhang M. Liquid-Liquid Phase Separation in Neuronal Development and Synaptic Signaling. Developmental Cell. 2020;55(1):18–29.

18. Ray S, Singh N, Pandey S, Kumar R, Gadhe L, Datta D, et al. Liquid-liquid phase separation and liquid-to-solid transition mediate alpha-synuclein amyloid fibril containing hydrogel formation. bioRxiv preprint. 2019:1–40.

19. Nedelsky NB, Taylor JP. Bridging biophysics and neurology: aberrant phase transitions in neurodegenerative disease. Nature Reviews Neurology. 2019;15(5):272–86.

20. Zhou Y, Su JM, Samuel CE, Ma D. Measles Virus Forms Inclusion Bodies with Properties of Liquid Organelles. Journal of Virology. 2019;93(21):e00948–19.

21. Seim I, Roden CA, Gladfelter AS. Role of spatial patterning of N-protein interactions in SARS-CoV-2 genome packaging. Biophysical Journal. 2021;120(14):2771–84.

22. Söding J, Zwicker D, Sohrabi-Jahromi S, Boehning M, Kirschbaum J. Mechanisms for Active Regulation of Biomolecular Condensates. Trends in Cell Biology. 2020;30(1):4–14.

23. Abyzov A, Blackledge M, Zweckstetter M. Conformational Dynamics of Intrinsically Disordered Proteins Regulate Biomolecular Condensate Chemistry. Chemical Reviews. 2022.

24. Biesaga M, Frigolé-Vivas M, Salvatella X. Intrinsically disordered proteins and biomolecular condensates as drug targets. Current Opinion in Chemical Biology. 2021;62:90–100.

25. Spanni S, Tereshchenko M, Mastromarco GJ, Ihn SJ, Lee HO. Biomolecular condensates in neurodegeneration and cancer. Traffic. 2019;20:890–911.

26. Jiang S, Fagman JB, Chen C, Alberti S, Liu B. Protein phase separation and its role in tumorigenesis. eLife. 2020;9:e60264.

27. Cai D, Liu Z, Lippincott-Schwartz J. Biomolecular Condensates and Their Links to Cancer Progression. Trends in Biochemical Sciences. 2021;46(7):535–49.

28. Taniue K, Akimitsu N. Aberrant phase separation and cancer. FEBS Journal. 2021.

29. Alberti S, Hyman AA. Are Aberrant Phase Transitions a Driver of Cellular Aging? Bioessays. 2016;38:959–68.

30. Peskett TR, Rau F, O’Driscoll J, Patani R, Lowe AR, Saibil HR. A Liquid To Solid Phase Transition Underlying Pathological Huntingtin Exon1 Aggregation. Mol Cell. 2018;70:588–601.

31. Riguet N, Mahul-Mellier A-L, Maharjan N, Burtscher J, Croisier M, Knott G, et al. Nuclear and cytoplasmic huntingtin inclusions exhibit distinct biochemical composition, interactome and ultrastructural properties. Nature Communications. 2021;12(1):6579.

32. Patel A, Lee HO, Jawerth L, Maharana S, Jahnel M, Hein MY, et al. A Liquid-to-Solid Phase Transition of the ALS Protein FUS Accelerated By Disease Mutation. Cell. 2015;162:1066–77.

33. Murthy AC, Dignon GL, Kan Y, Zerze GH, Parekh SH, Mittal J, et al. Molecular Interactions Underlying Liquid-Liquid Phase Separation of the FUS Low-Complexity Domain. Nature Structural and Molecular Biology. 2019;26:637–48.

34. Gibson BA, Doolittle LK, Schneider MWG, Jensen LE, Gamarra N, Henry L, et al. Organization of Chromatin by intrinsic and Regulated Phase Separation. Cell. 2019;179:470–84.

35. Shin Y, Chang Y-C, Lee DSW, Berry J, Sanders DW, Ronceray P, et al. Liquid Nuclear Condensates Mechanically Sense and Restructure the Genome. Cell. 2018;175:1481–91.

36. Mir M, Bickmore W, Furlong EEM, Narlika G. Chromatin topology, condensates and gene regulation: shifting paradigms or just a phase? Development. 2019;146(19):dev182766.

37. Dignon GL, Zheng W, Kim YC, Best RB, Mittal J. Sequence Determinants of Protein Phase Behavior from a Coarse-Grained Model. PLoS Computational Biology. 2018;14:e1005941.

38. Statt A, Casademunt H, Brangwynne CP, Panagiotopoulos AZ. Model for disordered proteins with strongly sequence-dependent liquid phase behaviour. J Chem Phys. 2020;152:075101.

39. Wang B, Zhang L, Dai T, Qin Z, Lu H, Zhang L, et al. Liquid–liquid phase separation in human health and diseases. Signal Transduction and Targeted Therapy. 2021;6(1):290.

40. Uversky VN. Intrinsically Disordered Proteins and Their Mysterious (Meta)Physics. Frontiers in Physics. 2019;7:10.

41. Peran I, Mittag T. Molecular structure in biomolecular condensates. Current Opinion in Structural Biology. 2020;60:17–26.

42. Bhandari K, Cotten MA, Kim J, Rosen MK, Schmit JD. Structure–Function Properties in Disordered Condensates. The Journal of Physical Chemistry B. 2021;125(1):467–76.

43. Orti F, Navarro AM, Rabinovich A, Wodak SJ, Marino-Buslje C. Insight into membraneless organelles and their associated proteins: Drivers, Clients and Regulators. Computational and Structural Biotechnology Journal. 2021;19:3964–77.

44. Choi J-M, Dar F, Pappu R. LASSI: A lattice model for simulating phase transitions of multivalent proteins. PLoS Computational Biology. 2019;15:e1007028.

45. Harmon TS, Holehouse AS, Rosen MK, Pappu RV. Intrinsically disordered linkers determine the interplay between phase separation and gelation in multivalent proteins. eLife. 2017;6:e30294.

46. Jafarinia H, Van der Giessen E, Onck PR. Phase Separation of Toxic Dipeptide Repeat Proteins Related to C9orf72 ALS/FTD. Biophysical Journal. 2020;119:843–51.

47. Shillcock JC, Brochut M, Chénais E, Ipsen JH. Phase behaviour and structure of a model biomolecular condensate. Soft Matter. 2020;16(27):6413–23.

48. Paloni M, Bailly R, Ciandrini L, Barducci A. Unraveling Molecular Interactions in Liquid-Liquid Phase Separation of Disordered Proteins by Atomistic Simulations. J Phys Chem B. 2020;124:9009–16.

49. Zheng W, Dignon GL, Jovic N, Xu X, Regy RM, Fawzi NL, et al. Molecular Details of Protein Condensates Probed by Microsecond Long Atomistic Simulations. J Phys Chem B. 2020;124:11671–9.

50. Dignon GL, Zheng W, Mittal J. Simulation methods for liquid-liquid phase separation of disorded proteins. Curr Op Chem Eng. 2019;23:92–8.

51. Hafner AE, Krausser J, Saric A. Minimal coarse-grained models for molecular self-organisation in biology. Curr Op Struct Biol. 2019;58:43–52.

52. Joseph JA, Reinhardt A, Aguirre A, Chew PY, Russell KO, Espinosa JR, et al. Physics-driven coarse-grained model for biomolecular phase separation with near-quantitative accuracy. Nature Computational Science. 2021;1(11):732–43.

53. Benayad Z, von Bülow S, Stelzl LS, Hummer G. Simulation of FUS Protein Condensates with an Adapted Coarse-Grained Model. Journal of Chemical Theory and Computation. 2021;17:525–37.

54. Tsanai M, Frederix Pim WJM, Schroer CFE, Souza PCT, Marrink SJ. Coacervate formation studied by explicit solvent coarse-grain molecular dynamics with the Martini model. Chemical Science. 2021.

55. Bari KJ, Prakashchand DD. Fundamental Challenges and Outlook in Simulating Liquid–Liquid Phase Separation of Intrinsically Disordered Proteins. The Journal of Physical Chemistry Letters. 2021;12(6):1644–56.

56. Joseph JA, Espinosa JR, Sanchez-Burgos I, Garaizar A, Frenkel D, Collepardo-Guevara R. Thermodynamics and kinetics of phase separation of protein-RNA mixtures by a minimal model. Biophysical Journal. 2021;120(7):1219–30.

57. Ranganathan S, Shakhnovich E. Effect of RNA on Morphology and Dynamics of Membraneless Organelles. The Journal of Physical Chemistry B. 2021;125(19):5035–44.

58. Tesei G, Schulze Thea K, Crehuet R, Lindorff-Larsen K. Accurate model of liquid–liquid phase behavior of intrinsically disordered proteins from optimization of single-chain properties. Proceedings of the National Academy of Sciences. 2021;118(44):e2111696118.

59. Ruff KM, Pappu RV, Holehouse AS. Conformational Preferences and Phase Behaviour of Intrinsically Disordered Low Complexity Sequences: Insights from Multiscale Simulations. Curr Op Struct Biol. 2019;56:1–10.

60. Wang J, Choi J-M, Holehouse AS, Lee HO, Zhang X, Jahnel M, et al. A Molecular Grammar Governing the Driving Forces for Phase Separation of Prion-like RNA Binding Proteins. Cell. 2018;174(3):688-99.e16.

61. Harmon TS, Holehouse AS, Pappu RV. Differential Solvation of Intrinsically Disordered Linkers Drives the Formation of Spatially Organised Droplets in Ternary Systems of Linear Multivalent Proteins. New J Physics. 2018;20:045002–16.

62. Rana U, Brangwynne CP, Panagiotopoulos AZ. Phase separation vs aggregation behavior for model disordered proteins. The Journal of Chemical Physics. 2021;155(12):125101.

63. Zhang Z, Chen Q, Colby RH. Dynamics of associative polymers. Soft Matter. 2018;14(16):2961–77.

64. Hoogerbrugge PJ, Koelman JMVA. Simulating Microscopic Hydrodynamic Phenomena with Dissipative Particle Dynamics. Europhys Lett. 1992;19:155–60.

65. Groot RD, Warren PB. Dissipative Particle Dynamics: Bridging the Gap Between Atomistic and Mesoscopic Simulations. J Chem Phys. 1997;107:4423–35.

66. Burke KA, Janke AM, Rhine CL, Fawzi NL. Residue-by-Residue View of In Vitro FUS Granules that Bind the C-Terminal Domain of RNA Polymerase II. Molecular Cell. 2015;60:231–41.

67. Ahlers J, Adams EM, Bader V, Pezzotti S, Winklhofer KF, Tatzelt J, et al. The key role of solvent in condensation: Mapping water in liquid-liquid phase-separated FUS. Biophysical Journal. 2021;120(7):1266–75.

68. Mitrea DM, Cika JA, Guy CS, Ban D, Banerjee PR, Stanley CB, et al. Nucleophosmin integrates within the nucleolus via multi-modal interactions with proteins displaying R-rich linear motifs and rRNA. eLife. 2016;5:e13571.

69. Barabási A-L. Network Science: Cambridge University Press; 2016 21 July 2016.

70. Newman MEJ. The Structure and Function of Complex Networks. SIAM Review. 2003;45(2):167–256.

71. Laniau J, Frioux C, Nicolas J, Baroukh C, Cortes MP, Got J, et al. Combining graph and flux-based structures to decipher phenotypic essential metabolites within metabolic networks. PeerJ. 2017;5:e3860.

72. Gheeraert A, Pacini L, Batista VS, Vuillon L, Lesieur C, Rivalta I. Exploring Allosteric Pathways of a V-Type Enzyme with Dynamical Perturbation Networks. J Phys Chem B. 2019;123(16):3452–61.

73. Dorantes-Gilardi R, Bourgeat L, Pacini L, Vuillon L, Lesieur C. In proteins, the structural responses of a position to mutation rely on the Goldilocks principle: not too many links, not too few. Phys Chem Chem Phys. 2018;20(39):25399–410.

74. Watts DJ, Strogatz SH. Collective dynamics of ‘small-world’ networks. Nature. 1998;393(6684):440–2.

75. Saramäki J, Kivelä M, Onnela J-P, Kaski K, Kertész J. Generalizations of the clustering coefficient to weighted complex networks. Physical Review E. 2007;75(2):027105.

76. Shillcock JC. OSPREY-DPD. 2020. p. Open Source Polymer Research Engine - Dissipative Particle Dynamics, https://github.com/Osprey-DPD/osprey-dpd.

77. Shillcock J, Lipowsky R. Equilibrium structure and lateral stress distribution of amphiphilic bilayers from dissipative particle dynamics simulations. Journal of Chemical Physics. 2002;117(10):5048–61.

78. Venturoli M, Sperotto MM, Kranenburg M, Smit B. Mesoscopic Models of Biological Membranes. Physics Reports. 2006;437:1–54.

79. Ortiz V, Nielsen S, Discher D, Klein M, Lipowsky R, Shillcock J. Dissipative particle dynamics simulations of polymersomes. Journal of Physical Chemistry B. 2005;109(37):17708–14.

80. Grafmueller A, Shillcock J, Lipowsky R. The Fusion of Membranes and Vesicles: Pathway and Energy Barriers from Dissipative Particle Dynamics. Biophysical Journal. 2009;96(7):2658–75.

81. Lipowsky R, Brinkmann M, Dimova R, Haluska C, Kierfeld J, Shillcock J. Wetting, budding, and fusion—morphological transitions of soft surfaces. Journal of Physics: Condensed Matter. 2005;17(31):S2885–S902.

82. Laradji M, Sunil Kumar PB. Domain growth, budding, and fission in phase-separating self-assembled fluid bilayers. The Journal of Chemical Physics. 2005;123(22):224902.

83. Espagnol P, Warren PB. Perspective: Dissipative Particle Dynamics. J Chem Phys. 2017;146:150901–17.

84. Ilnytskyi JM, Holovatch Y. How does the scaling for the polymer chain in the dissipative particle dynamics hold? Condensed Matter Physics. 2007;10:539–51.

85. Chen C, Ding X, Akram N, Xue S, Luo S-Z. Fuse in Sarcoma: Properties, Self-Assembly and Correlation with Neurodegenerative Diseases. Molecules. 2019;24:1622–39.

86. Murthy AC, Tang WS, Jovic N, Janke AM, Seo DH, Perdikari TM, et al. Molecular interactions contributing to FUS SYGQ LC-RGG phase separation and co-partitioning with RNA polymerase II heptads. Nature Structural & Molecular Biology. 2021;28(11):923–35.

87. Tomasso ME, Tarver MJ, Devarajan D, Whitten ST. Hydrodynamic radii of Intrinsically Disordered Proteins Determined from Experimental Polyproline II Propensities. PLoS Computational Biology. 2015;12:1–22.

88. Marsh JA, Forman-Kay JD. Sequence Determinants of Compaction in Intrinsically Disordered Proteins. Biophysical Journal. 2010;98:2383–90.

89. Rubinstein M, Colby RH. Polymer Physics. New York: Oxford University Press; 2003.

90. Dünweg B, Reith D, Steinhauser M, Kremer K. Corrections to scaling in the hydrodynamic properties of dilute polymer solutions. The Journal of Chemical Physics. 2002;117(2):914–24.

91. O’Flynn BG, Mittag T. The role of liquid–liquid phase separation in regulating enzyme activity. Current Opinion in Cell Biology. 2021;69:70–9.

92. Wang Y, Ghumare E, Vandenberghe R, Dupont P. Comparison of Different Generalizations of Clustering Coefficient and Local Efficiency for Weighted Undirected Graphs. Neural Comput. 2017;29(2):313–31.

93. Onnela J-P, Saramäki J, Kertész J, Kaski K. Intensity and coherence of motifs in weighted complex networks. Physical Review E. 2005;71(6):065103.

94. Elbaum-Garfinkle S, Kim Y, Szczepaniak K, Chen CC-H, Eckmann CR, Myong S, et al. The disordered P granule protein LAF-1 drives phase separation into droplets with tunable viscosity and dynamics. Proceedings of the National Academy of Sciences. 2015;112(23):7189.

95. Ronceray P, Zhang Y, Liu X, Wingreen NS. Stoichiometry Controls the Dynamics of Liquid Condensates of Associative Proteins. Physical Review Letters. 2022;128(3):038102.

96. Brangwynne CP, Tompa P, Pappu RV. Polymer Physics of Intracellular Phase Transitions. Nature Physics. 2015;11:899–904.

97. Rubinstein M, Dobrynin AV. Solutions of Associative Polymers. TRIP. 1997;5:181–6.

98. Zilman A, Tlusty T, Safran SA. Entropic networks in colloidal, polymeric and amphiphilic systems. Journal of Physics: Condensed Matter. 2002;15(1):S57–S64.

99. Dudowicz J, Freed KF. Lattice cluster theory of associating polymers. I. Solutions of linear telechelic polymer chains. The Journal of Chemical Physics. 2012;136(6):064902.

100. Holehouse AS, Pappu RV. Collapse Transitions of Proteins and the Interplay Among Backbone, Sidechain, and Solvent Interactions. Annu Rev Biophys. 2018;47:19–39.

101. Dobrynin AV. Phase Diagram of Solutions of Associative Polymers. Macromolecules. 2004;37(10):3881–93.

102. Bergsma J, Leermakers FAM, Kleijn JM, van der Gucht J. A Hybrid Monte Carlo Self-Consistent Field Model of Physical Gels of Telechelic Polymers. Journal of Chemical Theory and Computation. 2018;14(12):6532–43.

103. Arai N. Structural Analysis of Telechelic Polymer Solution Using Dissipative Particle Dynamics Simulations. Mol Sim. 2015;41:996–1001.

104. Manassero C, Raos G, Allegra G. Structure of Model Telechelic Polymer Melts by Computer Simulation. J Macromol Sci Part B: Physics. 2005;44:855–71.

105. Witten TA. Structured Fluids. Physics Today. 1990;43(7):21–8.

106. Rubinstein M, Dobrynin AV. Solutions of Associative Polymers. Trends in Polymer Science. 1997;5(6):181–6.

107. Bratek-Skicki A, Pancsa R, Meszaros B, Van Lindt J, Tompa P. A guide to regulation of the formation of biomolecular condensates. The FEBS Journal. 2020;287(10):1924–35.

108. Owen I, Rhoades S, Yee D, Wyne H, Gery K, Hannula I, et al. The prion-like domain of Fused in Sarcoma is phosphorylated by multiple kinases affecting liquid- and solid-liquid phase transitions. Molecular Biology of the Cell. 2020;31:2522–36.

109. Murakami T, Qamar S, Lin JQ, Schierle GSK, Rees E, Miyashita A, et al. ALS/FTD Mutation-Induced Phase Transition of FUS Liquid Droplets and Reversible Hydrogels into Irreversible Hydrogels Impairs RNP Granule Function. Neuron. 2015;88:678–90.

110. Jawerth L, Fischer-Friedrich E, Saha S, Wang J, Franzmann T, Zhang X, et al. Protein condensates as aging Maxwell fluids. Science. 2020;370(6522):1317–23.

111. Woodruff JB, Hyman AA, Boke E. Organization and Function of Non-dynamic Biomolecular Condensates. Trends in Biochemical Sciences. 2018;43(2):81–94.

112. Ray S, Singh N, Kumar R, Patel K, Pandey S, Datta D, et al. α-Synuclein aggregation nucleates through liquid–liquid phase separation. Nature Chemistry. 2020;12(8):705–16.

113. Risso-Ballester J, Galloux M, Cao J, Le Goffic R, Hontonnou F, Jobart-Malfait A, et al. A condensate-hardening drug blocks RSV replication in vivo. Nature. 2021;595:596–9.

114. Hofweber M, Dormann D. Friend or foe—Post-translational modifications as regulators of phase separation and RNP granule dynamics. Journal of Biological Chemistry. 2019;294(18):7137–50.

115. Perdikari TM, Jovic N, Dignon GL, Kim YC, Fawzi NL, Mittal J. A predictive coarse-grained model for position-specific effects of post-translational modifications. Biophysical Journal. 2021;120(7):1187–97.

116. Taylor NO, Wei MT, Stone HA, Brangwynne CP. Quantifying Dynamics in Phase-Separated Condensates Using Fluorescence Recovery after Photobleaching. Biophysical Journal. 2019;117(7):1285–300.

117. Marchetti-Bowick M, Yin J, Howrylak JA, Xing EP. A time-varying group sparse additive model for genome-wide association studies of dynamic complex traits. Bioinformatics. 2016;32(19):2903–10.

118. Latapy M, Viard T, Magnien C. Stream graphs and link streams for the modeling of interactions over time. Social Network Analysis and Mining. 2018;8(1):61.

119. Latapy M, Tabourier L, Armoux T. Predicting interactions between Individuals with structural and dynamical information. Journal of Interdisciplinary Methodologies and Issues in Sciences. 2019.

120. Qiu X, Zhao L, Wang J, Wang X, Wang Q. Effects of time-dependent diffusion behaviors on the rumor spreading in social networks. Physics Letters A. 2016;380(24):2054–63.

121. Moinet A, Starnini M, Pastor-Satorras R. Burstiness and Aging in Social Temporal Networks. Physical Review Letters. 2015;114(10):108701.

122. Idri A, Oukarfi M, Boulmakoul A, Zeitouni K, Masri A. A new time-dependent shortest path algorithm for multimodal transportation network. Procedia Computer Science. 2017;109:692–7.

123. Girase H, Gang H, Malla S, Li J, Kanehara A, Mangalam K, et al. LOKI: Long Term and Key Intentions for Trajectory Prediction. IEEE/CVF International Conference on Computer Vision; 11-17 October 2021; Online2021. p. 9803–12.

124. Bracha D, Walls MT, Brangwynne CP. Probing and engineering liquid-phase organelles. Nat Biotechnol. 2019;37(12):1435–45.

125. Elbaum-Garfinkle S, Kriwacki RW. Phase Separation in Biology & Disease: The next chapter. Journal of Molecular Biology. 2021;433(12):166990.

126. Hastings RL, Boeynaems S. Designer Condensates: A Toolkit for the Biomolecular Architect. Journal of Molecular Biology. 2021;433(12):166837.

127. Lohia R, Hansen M, Brannigan G. Contiguously hydrophobic sequences are functionally significant throughout the human exome. Biorxiv. 2021.

128. Garabedian MV, Wang W, Dabdoub JB, Tong M, Caldwell RM, Benman W, et al. Designer membraneless organelles sequester native factors for control of cell behavior. Nature Chemical Biology. 2021;17:998–1007.

129. Wheeler RJ, Therapeutics - how to treat phase separation-associated diseases. Emerging Topics in Life Sciences. 2020;4:331–42.

130. Humphrey W, Dalke A, Schulten K. VMD - Visual Molecular Dynamics. Journal of Molecular Graphics. 1996;14:33–8.

